# Confined Migration Induces Heterochromatin Formation and Alters Chromatin Accessibility

**DOI:** 10.1101/2021.09.22.461293

**Authors:** Chieh-Ren Hsia, Jawuanna McAllister, Ovais Hasan, Julius Judd, Seoyeon Lee, Richa Agrawal, Chao-Yuan Chang, Paul Soloway, Jan Lammerding

## Abstract

During migration, cells often squeeze through small constrictions, requiring extensive deformation. We hypothesized that the nuclear deformation associated with such confined migration could alter chromatin organization and function. Studying cells migrating through microfluidic devices that mimic interstitial spaces *in vivo*, we found that confined migration results in increased H3K9me3 and H3K27me3 heterochromatin marks that persist for several days. This “confined migration-induced heterochromatin” (CMiH) was distinct from heterochromatin formation during migration initiation. Confined migration predominantly decreased chromatin accessibility at intergenic regions near centromeres and telomeres, suggesting heterochromatin spreading from existing heterochromatin sites. Consistent with the overall decrease in chromatin accessibility, global transcription was decreased during confined migration. Intriguingly, we also identified increased accessibility at promoter regions of genes linked to chromatin silencing, tumor invasion, and DNA damage response. Inhibiting CMiH reduced migration speed, suggesting that CMiH promotes confined migration. Together, our findings indicate that confined migration induces chromatin changes that regulate cell migration and other cellular functions.

## Introduction

Cell migration is a crucial biological process required for many physiological functions (Luster et al., 2005; Scarpa and Mayor, 2016). Cell migration also plays a pivotal role in metastasis, which is the major cause of cancer-related death (Chaffer and Weinberg, 2011; Lambert et al., 2017). *In vivo*, cells must often squeeze through narrow interstitial spaces that are only 1-20 μm in diameter, and thus substantially smaller than the size of the cell (Kameritsch and Renkawitz, 2020; Weigelin et al., 2012). This recognition has led to an increased interest in studying cell migration in ‘confined’ environments, including the use of three-dimensional (3D) *in vitro* models, such as collagen matrices and microfluidic devices that resemble cell migration *in vivo* (Paul et al., 2017). Previous studies have revealed that the nucleus, which is the largest and most rigid organelle (Lammerding, 2011; Swift et al., 2013), undergoes severe deformation during confined migration (Denais et al., 2016; Irianto et al., 2016; Raab et al., 2016). The nuclear deformation and physical stress associated with confined migration can result in nuclear envelope rupture and cause DNA damage (Denais et al., 2016; Irianto et al., 2016; Raab et al., 2016; Shah et al., 2020). Furthermore, recent evidence suggests that migration through confined spaces can lead to rearrangements in 3D genome organization in neutrophils and cancer cells (Golloshi et al., 2020; Jacobson et al., 2018), which could affect their transcriptional regulation and cellular functions. However, the effect of confined migration on chromatin modifications and its functional consequences have not been explored.

Chromatin is found in two distinct states: the relaxed and transcriptionally active euchromatin, and the condensed and transcriptionally silenced heterochromatin (Janssen et al., 2018; Strålfors and Ekwall, 2011). These states are controlled by post-translational histone modifications (Bannister and Kouzarides, 2011) and DNA methylation (Rose and Klose, 2014). For example, trimethylation of lysine 9 on histone H3 (H3K9me3) is associated with constitutive heterochromatin, which mostly locates in centromeric and telomeric DNA regions (Janssen et al., 2018; Strålfors and Ekwall, 2011; Canzio et al., 2011); trimethylation of lysine 27 on histone H3 (H3K27me3) is associated with facultative heterochromatin, which is developmentally-regulated (Janssen et al., 2018; Strålfors and Ekwall, 2011; Wu et al., 2016). In contrast, acetylation of lysine 9 on histone H3 (H3K9ac) is associated with euchromatin and active promoters (Lennartsson and Ekwall, 2009). Increased H3K9me3 and H3K27me3 chromatin modifications are crucial to initiate 2D cell migration and for transwell migration (Gerlitz, 2020; Gerlitz and Bustin, 2010; Liu et al., 2018; B. Zhang et al., 2016; X. Zhang et al., 2016), potentially by repression of specific genes (Segal et al., 2018). At the same time, increased euchromatin facilitates cell migration through 3D collagen matrices (Fischer et al., 2020; Wang et al., 2018), likely by increasing nuclear deformability (Fischer et al., 2020; Stephens et al., 2018). Intriguingly, recent studies have found that the physical microenvironment of cells can cause changes in chromatin modifications: mechanical compression of cells induces reversible chromatin condensation with increased H3K9me3 and H3K27me3 marks (Damodaran et al., 2018), whereas cyclic stretching of cells leads to rapid and transient loss of heterochromatin (Nava et al., 2020). Therefore, we aimed to explore whether nuclear deformation during confined migration could result in altered chromatin modifications, genomic accessibility, and transcriptional activity, and whether such changes could modulate the ability of cells to migrate through confined 3D environments.

We found that confined 3D migration in microfluidic devices induced persistent global heterochromatin formation in cancer cells and fibroblasts, well beyond the changes observed during unconfined migration. The increased heterochromatin formation increased with the degree of confinement, depended on histone modifying enzymes, and was modulated by nuclear envelope proteins and stretch-sensitive ion channels. Although we did not observe global changes of heterochromatin levels in cells migrating in 3D collagen matrices with variable pore sizes, Assay for Transposase-Accessible Chromatin using sequencing (ATAC-seq) analysis revealed that cell migration in collagen matrices with smaller pore sizes led to reduced chromatin accessibility at intergenic regions near centromeres and telomeres, consistent with the increased heterochromatin formation observed in the microfluidic devices. Global reduction of transcriptional activity after confined migration further supports the predominantly repressing accessibility change. On the other hand, we identified increased chromatin accessibility at promoter regions of genes responsible for a wide range of pathways, such as chromatin silencing, tumor invasion, and DNA damage response. Preventing heterochromatin formation using histone methyltransferase inhibitors resulted in impaired migration in the microfluidic devices, particularly through small constrictions, suggesting that confined migration-induced heterochromatin promotes confined migration.

## Results

### Heterochromatin increases during and after cell migration through confined microfluidic channels

To investigate the effect of confined cell migration and the associated nuclear deformation on histone modifications, we studied cells migrating through custom-made polydimethylsiloxane (PDMS) microfluidic devices with precisely defined constrictions that mimic interstitial spaces (Fig. 1A) (Davidson et al., 2015; Denais et al., 2016). These microfluidic devices enable us to control the pore sizes that cells encounter during 3D migration independent of the extracellular matrix concentration and stiffness. HT1080 fibrosarcoma cells migrated either through channels containing three rows of small constrictions (≤2×5 μm^2^ in cross-section) that require substantial nuclear deformation to squeeze through (“confined channels”), or through larger “control channels” (15×5 μm^2^ in cross-section) that do not require substantial nuclear deformation for transit. Cells were categorized according to their locations in the device: (1) in the unconfined area “before” entering the channels, (2) in the middle of “squeezing” through the confined constrictions (or passing through the larger control constrictions), or (3) “after” migration out of the channels into the unconfined area (Fig. 1A). For a detailed break-down, we also defined different zones within the 5-µm tall sections: “Zone 1” encompassed cells that that passed through the first row of constrictions but had not yet entered the second row; “Zone 2” defined cells in the area between the second and third row of constrictions (Fig. 1A). Cells were subsequently fixed and immunofluorescently labeled for heterochromatin marks (H3K27me3 and H3K9me3) and a euchromatin mark (H3K9ac) (Fig. 1B). To distinguish between true changes in chromatin modifications and effects of physical compression of the nuclear content due to deformation, we normalized the heterochromatin mark intensity to the euchromatin mark intensity in each cell. Increased heterochromatin formation should result in an increased ratio of heterochromatin marks to euchromatin marks, whereas physical compression of chromatin would increase both marks, and thus not alter their ratio. This ratio of heterochromatin marks to euchromatin marks represents the normalized heterochromatin level, which we refer to as “heterochromatin level” throughout the article.

**Figure 1:**
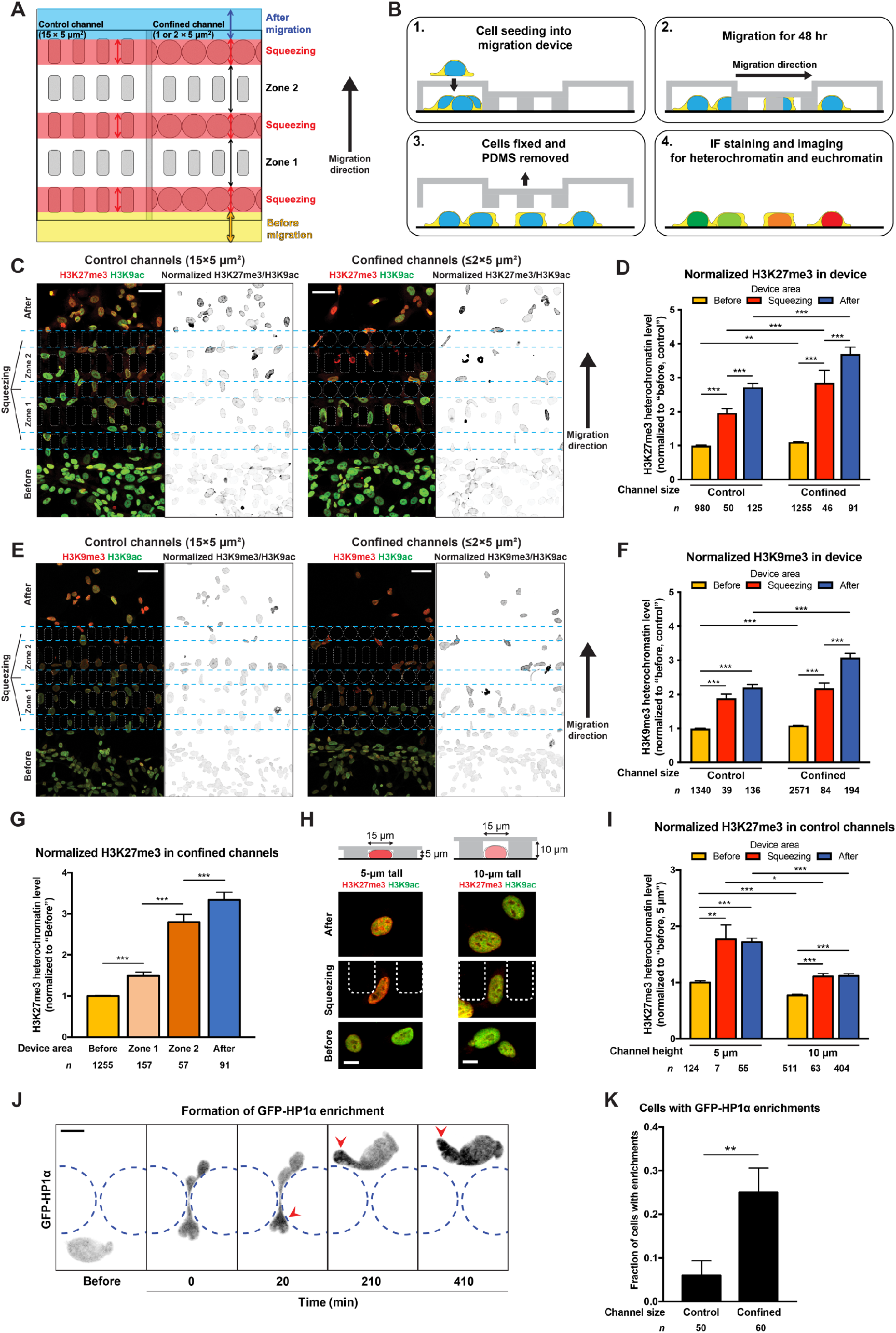
Confined migration induces heterochromatin formation in microfluidic migration devices. **(A)** A simplified schematic illustrating the design and the different areas in the migration device. Yellow indicates the unconfined area “before” cells entering the device. Red indicates the area while cells are “squeezing” or passing through the constrictions. Blue indicates the unconfined area “after” cells migrating out of the channels. “Zone 1” indicates the area after cells made one nuclear transit through the first row of constrictions. “Zone 2” indicates the area after cells made two nuclear transits through the first two rows of constrictions. **(B)** The workflow of staining experiments using PDMS microfluidic migration devices. Unlabeled cells were seeded into the device (step 1). Cells were allowed migration for 48 hours (step 2) before fixation and removal of the PDMS (step 3), and the cells were stained for chromatin marks (step 4). **(C)** Representative staining of H3K27me3 (red) and H3K9ac (green) in HT1080 cells migrating in a migration device, in control or confined channels. Normalized heterochromatin level is shown in inverted grayscale. Scale bars: 40 μm. **(D)** Quantification of normalized H3K27me3 heterochromatin level in HT1080 cells migrating in migration devices. All values are normalized to control channels “before” cells. \*\**p* < 0.01, \*\*\**p* < 0.001, two-way ANOVA with Tukey’s multiple comparison test. **(E)** Representative staining of H3K9me3 (red) and H3K9ac (green) in HT1080 cells migrating in a migration device, in control or confined channels. Normalized heterochromatin level is shown in inverted grayscale. Scale bars: 40 μm. **(F)** Quantification of normalized H3K9me3 heterochromatin level in HT1080 cells migrating in migration devices. All values are normalized to control channels “before” cells. \*\*\**p* < 0.001, two-way ANOVA with Tukey’s multiple comparison test. **(G)** Comparison between normalized H3K27me3 heterochromatin level of HT1080 cells in each area of the device. All values are normalized to “before” cells. \*\*\**p* < 0.001, one-way ANOVA with Tukey’s multiple comparison test. **(H)** Representative staining of H3K27me3 (red) and H3K9ac (green) in HT1080 cells migrating in devices with 5 μm or 10 μm-tall control channel. Cross-section view of channel design is shown above. Scale bars: 10 μm. **(I)** Quantification of normalized H3K27me3 heterochromatin level in HT1080 cells migrating in devices with 5-μm or 10-μm tall control channels. All values are normalized to 5 μm-tall control channels “before” cells. \**p* < 0.05, \*\**p* < 0.01, \*\*\**p* < 0.001, two-way ANOVA with Tukey’s multiple comparison test. **(J)** Representative inverted grayscale image sequence of local enrichment formation of GFP-HP1α (arrowhead) in HT1080 cells. Scale bar: 10 μm. Please see also Video S1. **(K)** Quantification of HT1080 cells with local enrichment of GFP-HP1α in control and confined channels. \*\**p* < 0.01, student’s *t* test with Welch’s correction for unequal variances. All values are normalized to “before” cells. \*\*\**p* < 0.001, one-way ANOVA with Tukey’s multiple comparison test. Data are presented as mean ± SEM, based on *n* cells (listed in each graph) pooled from three biological replicates, except for data in (I), which are based on two biological replicates.

Migration of HT1080 cells resulted in increased H3K27me3 heterochromatin levels as the cells squeezed through the ≤2×5 μm^2^ confined constrictions, and an even larger increase in cells that had exited the confined channels, compared to cells before entering the channels (Fig. 1C, D; Fig. S1A). Staining for H3K9me3 showed similar results, although the facultative heterochromatin (H3K27me3) levels increased more substantially as cells migrated through the confined channels (Fig. 1E, F; Fig. S1B). Migration of skin fibroblasts and MDA-MB-231 breast cancer cells through the device yielded similar results (Fig. S2A-F). Cells exhibited a progressive increase of heterochromatin levels as they migrated through the device (Fig. 1G; Fig. S1C ; Fig. S2G), whereas the intensity of the H3K9ac euchromatin mark remained constant or even decreased (Fig. S2H-M). The progressive increase in heterochromatin marks, along with the fact that we did not observe cells with the same high levels of normalized H3K27m3 marks in the “before” population as seen in the “after” population (Fig. S1C), supports the interpretation that cells accumulate heterochromatin marks in response to confined migration, instead of an enrichment of a pre-existing sub-population with high heterochromatin levels. Nonetheless, we cannot exclude the possibility that selection further contributes to the observed enrichment in heterochromatin marks. Since heterochromatin-associated histone modifications such as H3K9me3 can lead to the recruitment of DNA methylation machinery (Rose and Klose, 2014), we hypothesized that the increased heterochromatin histone marks would coincide with an increase in DNA methylation following migration through the devices. Indeed, migration through both control and confined channels increased 5-methylcytosine (5-meC) intensities (Fig. S1N, O).

Surprisingly, even cells migrating through the 15×5 μm^2^ control channels displayed an increase in heterochromatin levels. Although the effect was less pronounced than in the ≤2×5 μm^2^ confined channels (Fig. 1C-F), it suggests that the vertical confinement of 5 μm is sufficient to induce a nuclear response, consistent with recent observations that cell confinement to heights below 5 to 7 μm is sufficient to trigger mechanosensing responses of the nucleus (Lomakin et al., 2020; Venturini et al., 2020). To determine whether the observed increase in heterochromatin was due to physical confinement or merely represented chromatin changes associated with migration initiation (Gerlitz and Bustin, 2010), we conducted experiments in which cells migrated either through the 15×5 μm^2^ control channels or through taller (15×10 μm^2^) channels that do not result in vertical confinement of the nucleus (Fig. 1H). Cells migrating through the 5-μm tall channels exhibited a significantly larger increase in heterochromatin than cells migrating through the 10-μm tall channels (Fig. 1H, I), demonstrating that the observed effect is primarily attributed to the confinement and not the migration process *per se*. Collectively, these data suggest that confined migration can result in increased and persistent heterochromatin formation, which we termed “confined migration-induced heterochromatin” (CMiH).

### Heterochromatin formation is induced after nuclear transit

Since immunofluorescence staining is limited to end-point measurements, we performed time-lapse experiments with cells stably expressing GFP-HP1α, a heterochromatin reporter that directly binds to H3K9me3 but also requires the cooperative interaction of H3K27me3 for stable binding (Boros et al., 2014; Cheutin et al., 2003). We frequently observed local and persistent enrichments of GFP-HP1α in nuclei during cell migration through confined channels compared to control channels (Fig. 1J, K; Fig. S3A, B). The GFP-HP1α enrichments lasted for more than four hours after nuclear transit through the confined constrictions (Fig. 1J; Fig. S3A, b; Video S1), suggesting that the increase in GFP-HP1α signal reflects persistent chromatin modifications and not transient pooling of mobile GFP-HP1α. To further validate that the confined migration-induced GFP-HP1α enrichment reflects true heterochromatin formation, we performed fluorescence recovery after photobleaching (FRAP) analysis (Schmiedeberg et al., 2004) on the GFP-HP1α enriched nuclear regions, with centromeric GFP-HP1α foci serving as positive controls (Fig. S3C-H). Nuclear blebs with enriched GFP-HP1α signals had a decreased fraction of mobile GFP-HP1α compared to non-bleb nuclear regions (Fig. S3G, H), suggesting that these enrichments reflect GFP-HP1α bound to heterochromatin formed in nuclear bleb regions.

### Confined migration-induced heterochromatin formation is persistent

To assess the persistence of CMiH, we performed long-term time-lapse imaging of cells in the migration device in an incubator-mounted microscope combined with subsequent immunofluorescence analysis for H3K27me3 and H3K9ac histone marks (Fig. 2A; Video S2), enabling us to correlate heterochromatin levels with the time after nuclear transit through the constrictions for individual cells. Intriguingly, the heterochromatin levels in cells that had passed through the confined channels remained significantly elevated compared to cells in the unconfined “before” area during the 5-day observation period (Fig. 2B). Cells maintained CMiH even after completing at least one round of mitosis, without any trend of reversion in their heterochromatin levels (Fig. 2C; Fig. S4A, B), suggesting that the epigenetic modifications were inheritable through DNA replications. Cells that did not undergo mitosis after nuclear transit even showed a trend towards further increase in heterochromatin levels overtime (Fig. 2D; Fig. S4A, C). Overall, these experiments indicate that CMiH persists for at least 5 days, including in proliferating cells.

**Figure 2:**
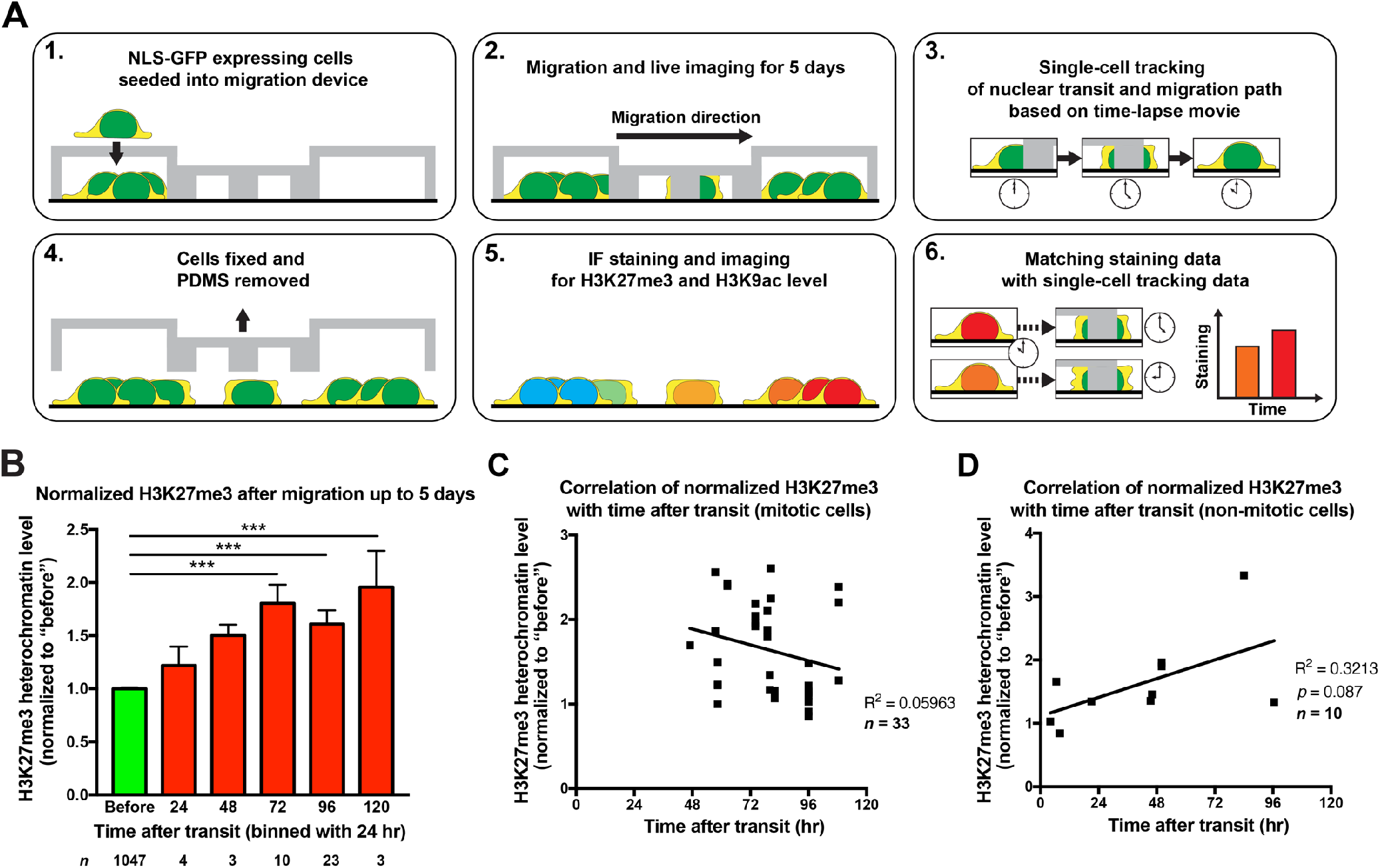
Confined migration induces persistent heterochromatin formation lasting for at least 5 days. **(A)** The workflow of live imaging with staining experiments using the migration devices. NLS-GFP expressing HT1080 cells were seeded into the device (step 1). Cells were allowed migration for up to 5 days under live imaging (step 2). The acquired time-lapse movie was then used to track single-cell migration path during the experiment, especially the timing of the last nuclear transit through confined constrictions (step 3). The cells were fixed, and the PDMS was removed (step 4), and the cells were then stained for chromatin marks (step 5). Finally, single-cell staining data were matched to the time-lapse tracking data for the correlation of heterochromatin level with the time after nuclear transit (step 6). Please see also Video S2. **(B)** The correlation of normalized H3K27me3 heterochromatin level with time after the last nuclear transit through confined constrictions in HT1080 cells, binned by an interval of 24 hours. All values are normalized to “before” cells. \*\*\**p* < 0.001, one-way ANOVA with Tukey’s multiple comparison test. **(C)** The correlation and linear regression of normalized H3K27me3 heterochromatin level with time after the last nuclear transit through confined constrictions in HT1080 cells with mitosis. All values are normalized to “before” cells. **(D)** The correlation and linear regression of normalized H3K27me3 heterochromatin level with time after the last nuclear transit through confined constrictions in HT1080 cells without any mitosis. All values are normalized to “before” cells. Data are presented as mean ± SEM, based on *n* cells (listed in each graph) pooled from three biological replicates.

### Confined migration-induced heterochromatin formation is dependent on histone modifying enzymes

To investigate the molecular mechanisms behind CMiH formation, we inhibited specific steps of the euchromatin-heterochromatin transition (Fig. 3A). Histone methyltransferases are required for the addition of methyl groups to H3K9 and H3K27. Treatment with 3-Deazaneplanocin A (DZNep), a broadband histone methyltransferase inhibitor (HMTi) (Miranda et al., 2009; Stephens et al., 2018) (Fig. S5A), significantly reduced CMiH (Fig. 3B), confirming that the observed changes in H3K27me3 and H3K9me3 heterochromatin levels were the result of increased histone methylation. We used a broadband histone deacetylase inhibitor (pan-HDACi), Trichostatin A (TSA) (Vigushin et al., 2001), to block the removal of histone acetylation (Fig. S5A), a pre-requisite for methylation of the same residues. Pan-HDACi treatment significantly reduced CMiH compared to vehicle controls (Fig. 3C), suggesting that the process is dependent on HDACs. One particular HDAC family member, HDAC3, had previously been implicated in catalyzing heterochromatin formation when cells are subjected to external compression or constraint spreading (Alisafaei et al., 2019; Damodaran et al., 2018). Thus, we hypothesized that HDAC3 may also play a crucial role in CMiH formation. Indeed, treatment with RGFP966, a potent HDAC3 inhibitor (HDAC3i) (Malvaez et al., 2013, p. 3) (Fig. S5B), demonstrated nearly the same inhibitory effect on CMiH as pan-HDACi (Fig. 3d). Depletion of HDAC3 by siRNA (Fig. S5C) similarly reduced CMiH after migration through confined channels, compared to non-target siRNA control (Fig. S5D). Combined treatment of HMTi and pan-HDACi completely eliminated CMiH, while also reducing the baseline heterochromatin levels before migration (Fig. 3E), speaking to the pivotal role of HMTs and HDACs in chromatin modifications. To test if the transition from heterochromatin back to euchromatin by histone demethylases (HDMs) also affects the levels of CMiH, we used JIB-04, a Jumonji-domain histone demethylases inhibitor (HDMi) (Parrish et al., 2018) (Fig. S5E). As expected, HDMi treatment increased CMiH in cells squeezing through constrictions and after migrating out of the channels, compared to vehicle controls (Fig. 3F).

**Figure 3:**
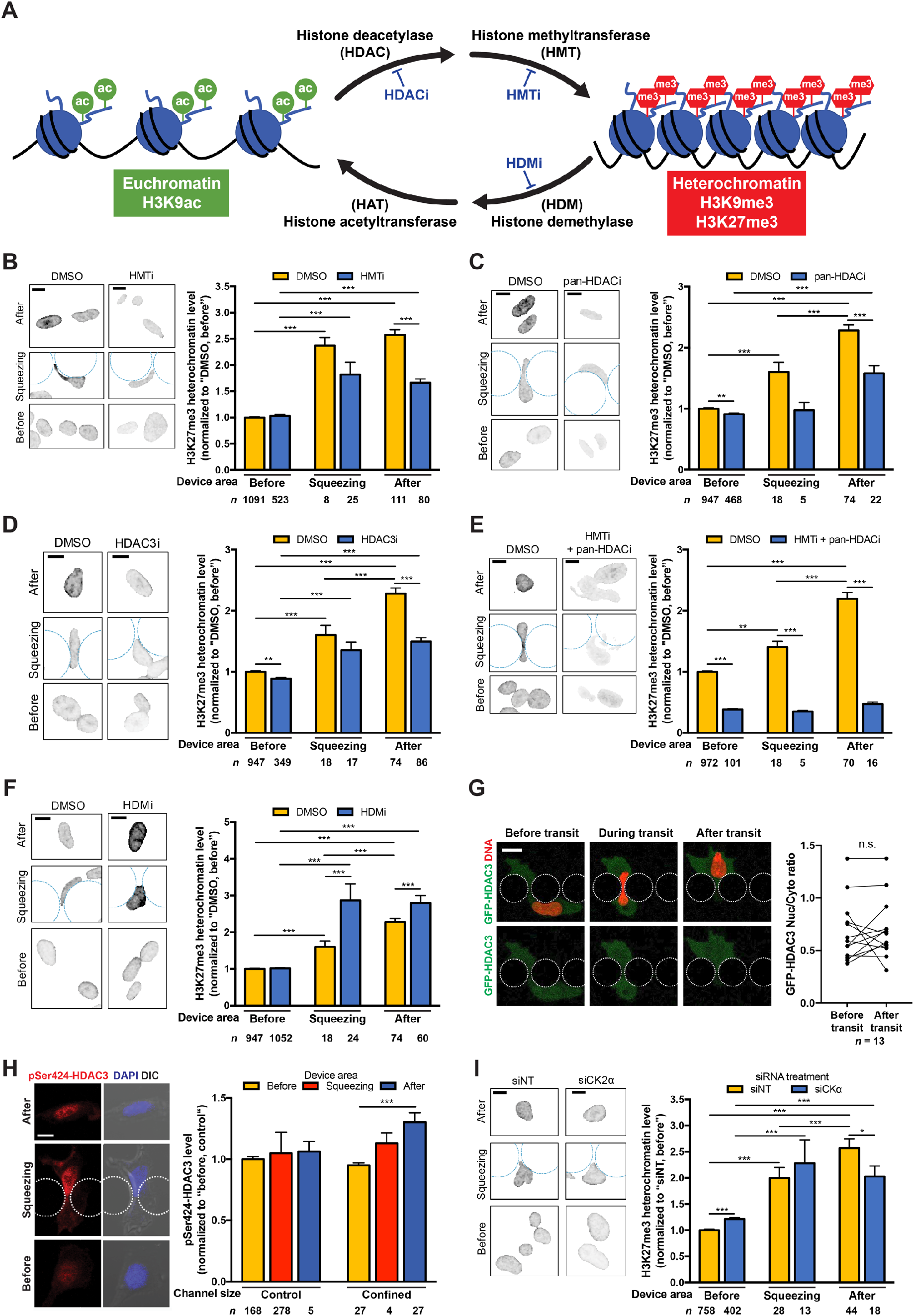
Confined migration-induced heterochromatin formation is dependent on histone modifying enzymes. **(A)** A simplified schematic of the histone modifying enzymes and the corresponding inhibitors involved in the transition between euchromatin state (left, in green) and heterochromatin state (right, in red). **(B)** Left panel: Representative images of normalized H3K27me3 heterochromatin in HT1080 cells treated with DMSO (vehicle) control or HMTi during confined migration. Scale bars: 10 μm. Right panel: Quantification of normalized H3K27me3 heterochromatin in cells treated with DMSO (vehicle) control or HMTi during confined migration. All values are normalized to DMSO “before” cells. \*\*\**p* < 0.001, two-way ANOVA with Tukey’s multiple comparison test. **(C)** Left panel: Representative images of normalized H3K27me3 heterochromatin in HT1080 cells treated with DMSO (vehicle) control or pan-HDACi during confined migration. Scale bars: 10 μm. Right panel: Quantification of normalized H3K27me3 heterochromatin in cells treated with DMSO (vehicle) control or pan-HDACi during confined migration. All values are normalized to DMSO “before” cells. \*\**p* < 0.01, \*\*\**p* < 0.001, two-way ANOVA with Tukey’s multiple comparison test. **(D)** Left panel: Representative images of normalized H3K27me3 heterochromatin in HT1080 cells treated with DMSO (vehicle) control or HDAC3i during confined migration. Scale bars: 10 μm. Right panel: Quantification of normalized H3K27me3 heterochromatin in cells treated with DMSO (vehicle) control or HDAC3i during confined migration. All values are normalized to DMSO “before” cells. \*\**p* < 0.01, \*\*\**p* < 0.001, two-way ANOVA with Tukey’s multiple comparison test. **(E)** Left panel: Representative images of normalized H3K27me3 heterochromatin in HT1080 cells treated with DMSO (vehicle) control or HMTi + pan-HDACi during confined migration. Scale bars: 10 μm. Right panel: Quantification of normalized H3K27me3 heterochromatin in cells treated with DMSO (vehicle) control or HMTi + pan-HDACi during confined migration. All values are normalized to DMSO “before” cells. \*\**p* < 0.01, \*\*\**p* < 0.001, two-way ANOVA with Tukey’s multiple comparison test. **(F)** Left panel: Representative images of normalized H3K27me3 heterochromatin in HT1080 cells treated with DMSO (vehicle) control or HDMi during confined migration. Scale bars: 10 μm. Right panel: Quantification of normalized H3K27me3 heterochromatin in cells treated with DMSO (vehicle) control or HDMi during confined migration. All values are normalized to DMSO “before” cells. \*\*\**p* < 0.001, two-way ANOVA with Tukey’s multiple comparison test. **(G)** Left panel: Representative image sequences of an HT1080 cell expressing GFP-HDAC3 (green) and stained with SPY555-DNA (red) migrating through a confined constriction. Scale bar: 20 μm. Right panel: Quantification of GFP-HDAC3 nucleoplasmic-to-cytoplasmic (Nuc/Cyto) ratio changes within an hour of before and after nuclear transit. N.S. not significant based on paired *t* test (*p* = 0.531). Please see also Video S3. **(H)** Left panel: Representative images of pSer424-HDAC3 (red) and DAPI (blue) in HT1080 cells during migration through control and confined channels. Scale bars: 10 μm. Right panel: Quantification of pSer424-HDAC3 intensities during confined migration. All values are normalized to control channels “before” cells. \*\*\**p* < 0.001, two-way ANOVA with Tukey’s multiple comparison test. **(I)** Left panel: Representative images of normalized H3K27me3 heterochromatin in HT1080 cells treated with non-target siRNA (siNT) or CK2α siRNA (siCK2α) during confined migration. Scale bar: 10 μm. Right panel: Quantification of normalized H3K27me3 heterochromatin in cells treated with siNT or siCK2α during confined migration. All values are normalized to siNT “before” cells. \**p* < 0.05, \*\*\**p* < 0.001, two-way ANOVA with Tukey’s multiple comparison test. Data are presented as mean ± SEM, based on *n* cells (listed in each graph) pooled from three biological replicates, except for data in (G), which are based on two biological replicates.

### Confined migration-induced heterochromatin formation is controlled by HDAC3 phosphorylation

To investigate the molecular mechanism responsible for CMiH in more detail, we examined the intracellular localization of HDAC3 during confined migration, as previous studies had reported that mechanical stimulation can trigger cytoplasmic to nuclear translocation of HDAC3 to induce heterochromatin formation (Alisafaei et al., 2019; Damodaran et al., 2018). However, live cell microscopy of cells stably expressing GFP-HDAC3 did not reveal any changes in the nucleoplasmic-to-cytoplasmic (Nuc/Cyto) ratio of GFP-HDAC3 after nuclear transit through confined constrictions (Fig. 3G; Video S3), suggesting that nuclear translocation of HDAC3 does not play a major role in CMiH formation. An alternative hypothesis is that confined migration results in increased activation of HDAC3, thereby contributing to increased heterochromatin formation. To test this hypothesis, we stained for phosphorylation of HDAC3 on Serine 424 (pSer424-HDAC3), which is critical for HDAC3 enzymatic activity (Zhang et al., 2005). Cells migration through confined channels, but not the larger control channels, led to increased levels of pSer424-HDAC3 (Fig. 3H). Since this particular HDAC3 phosphorylation is catalyzed by Casein Kinase 2 (CK2) (Zhang et al., 2005), we tested whether depletion of CK2α, a key subunit of CK2 (Litchfield, 2003, p. 2) (Fig. S5F), could inhibit CMiH formation. We observed reduced CMiH after cells depleted of CK2α migrated through confined channels, when compared to non-target siRNA control (Fig. 3I). However, the effect was mild and was not mirrored in cells squeezing through constrictions. This suggests that either residual CK2 levels are sufficient for CMiH, or that additional, yet to be determined mechanisms regulating HDAC3 activity are involved in CMiH formation.

### Stretch-sensitive ion channels, but not nuclear envelope rupture, contribute to confined migration-induced heterochromatin formation

To identify additional molecular players involved in CMiH formation, we investigated the role of the nuclear envelope (NE) proteins emerin and lamin A/C. HDAC3 localization to the nuclear periphery plays a crucial role in maintaining repression for specific gene loci during differentiation (Poleshko et al., 2017), and HDAC3 binding to the NE protein emerin activates HDAC3’s enzymatic activity and promotes its localization to the nuclear lamina (Demmerle et al., 2012). However, depletion of emerin by siRNA (Fig. S6A) did not reduce CMiH after migration, but instead resulted in increased heterochromatin levels as cells were squeezing through the constrictions (Fig. S6B). Similarly, depletion of the NE proteins lamin A/C (Fig. S6C), which play important roles in chromatin organization and help recruit emerin to the inner nuclear membrane (Poleshko et al., 2017; van Steensel and Belmont, 2017), did not reduce CMiH but increased the heterochromatin levels during severe nuclear deformation of cell squeezing (Fig. S6D). As lamin A/C and emerin have pleiotropic effects on chromatin organization and dynamics (Bronshtein et al., 2015; Dechat et al., 2009; Ranade et al., 2019), the exact mechanisms by which lamin A/C and emerin affect heterochromatin levels during cell squeezing remain to be explored.

Recent studies found that stretch-sensitive ion channels can modulate heterochromatin changes in response to mechanical challenges (Nava et al., 2020; Stephens et al., 2018). To test if CMiH is dependent on stretch-sensitive ion channels, we treated cells with gadolinium chloride (GdCl_3_), a broad inhibitor for stretch-sensitive ion channels (Caldwell et al., 1998). GdCl_3_-treated cells had significantly reduced CMiH compared to vehicle treated controls (Fig. 4A), indicating that stretch-sensitive ion channels contribute to CMiH, for example, via influx of extracellular calcium and/or the release of intracellular calcium from the nuclear lumen and endoplasmic reticulum (ER) (Lomakin et al., 2020; Venturini et al., 2020). A related mechanism may be the transient loss of NE integrity, termed “NE rupture”, which is due to the physical stress on the nucleus during confined migration (Denais et al., 2016; Irianto et al., 2016; Raab et al., 2016). NE rupture could result in calcium release from the nuclear lumen and ER, as well as mislocalization of histone modifying enzymes, ions, or signaling molecules (Denais et al., 2016; Irianto et al., 2016; Raab et al., 2016), thereby leading to heterochromatin formation. To investigate the role of NE rupture in CMiH, we performed time-lapse migration experiments of cells expressing a fluorescent reporter (cGAS-mCherry) that accumulates at the site of NE rupture (Denais et al., 2016), and then fixed the cells to stain for histone marks (Fig. 4B). Notably, we did not detect any significant difference in CMiH between cells that incurred NE rupture during migration and those that did not (Fig. 4C; Fig. S6E). Furthermore, cells that had experienced NE rupture did not show any significant correlation between the time that had passed since rupture and their heterochromatin levels (Fig. 4D; Fig. S6F). Taken together, these data suggest that CMiH is independent of NE rupture.

**Figure 4:**
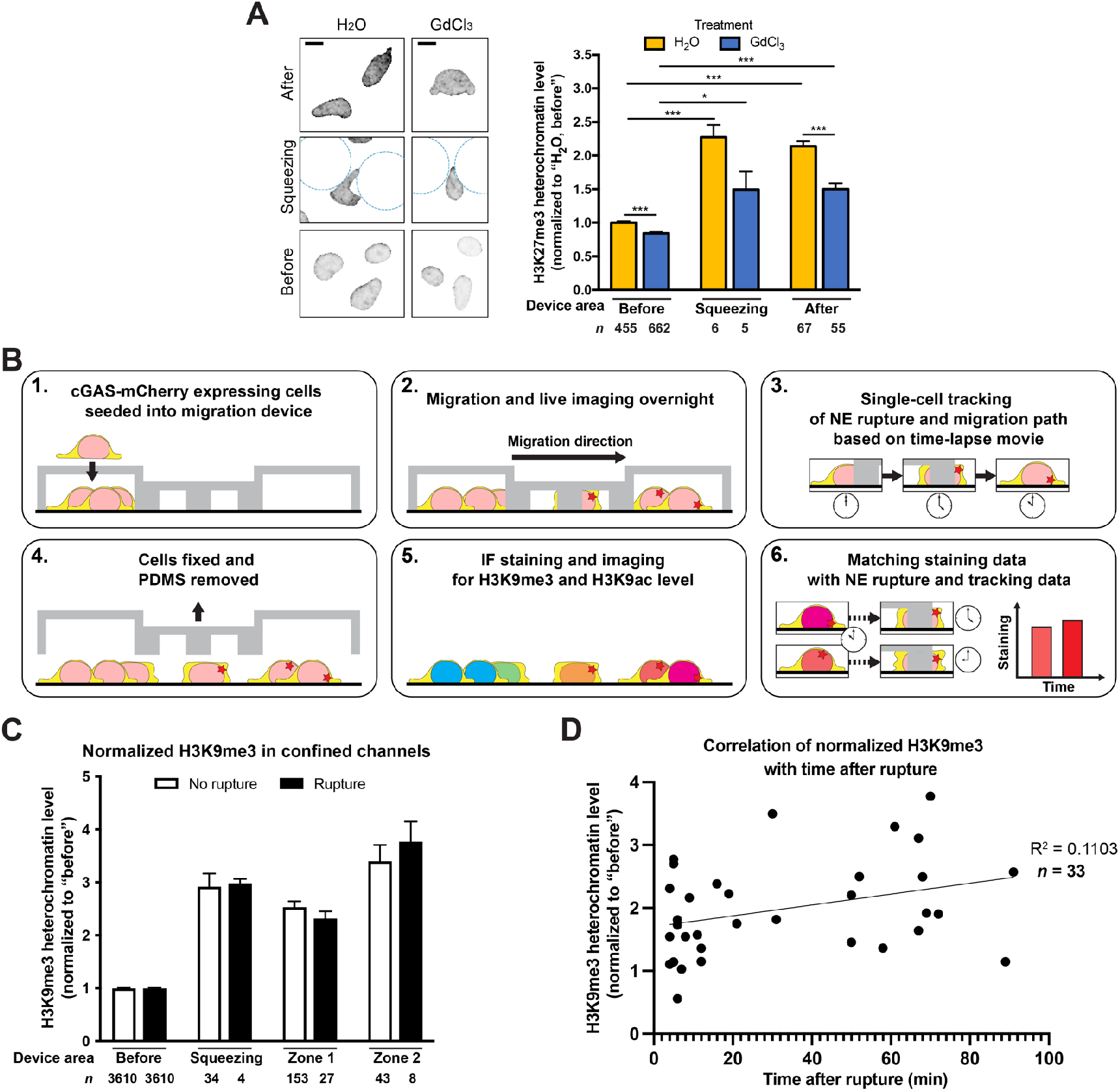
Confined migration-induced heterochromatin formation is affected by stretch-sensitive ion channels, but independent of nuclear envelope rupture. **(A)** Left panel: Representative images of normalized H3K27me3 heterochromatin in HT1080 cells treated with H_2_O (vehicle) control or GdCl_3_ during confined migration. Scale bars: 10 μm. Right panel: Quantification of normalized H3K27me3 heterochromatin in cells treated with vehicle (H_2_O) control or GdCl_3_ during confined migration. All values are normalized to H_2_O “before” cells. \**p* < 0.05, \*\*\**p* < 0.001, two-way ANOVA with Tukey’s multiple comparison test. **(B)** The workflow of live imaging with staining experiments using the migration devices. cGAS-mCherry expressing cells were seeded into the device (step 1). Cells were allowed migration overnight under live imaging (step 2). The acquired time-lapse movie was then used to track single-cell migration path during the experiment, especially the timing of the nuclear envelope (NE) rupture, marked by cGAS-mCherry foci formation (step 3). The cells were fixed, and the PDMS was removed (step 4), and the cells were then stained for chromatin marks (step 5). Finally, single-cell staining data were matched to the time-lapse tracking data for the correlation of heterochromatin level with the time after NE rupture (step 6). **(C)** Quantification of normalized H3K9me3 heterochromatin levels in HT1080 cells experienced NE rupture (orange) or not (blue) during confined migration. All “rupture” and “no rupture” heterochromatin levels are normalized the same level in “before” cells. **(D)** Correlation and linear regression of normalized H3K9me3 heterochromatin levels with time after rupture in HT1080 cells. All values are normalized to “before” cells. Data are presented as mean ± SEM, based on *n* cells (listed in each graph) pooled from three biological replicates, except for data in (C) and (D), which are based on two biological replicates.

### Confined migration alters chromatin accessibility

Histone modifications often result in altered chromatin accessibility and gene expression. Increased chromatin accessibility near promoters allows binding of transcriptional machineries and regulatory elements, which can lead to increased gene expression (Buenrostro et al., 2015; Tsompana and Buck, 2014). In contrast, decreased chromatin accessibility is associated with silenced genomic regions and heterochromatin (Buenrostro et al., 2015; Tsompana and Buck, 2014). Since the limited number (< 100) of cells that each microfluidic migration device can provide is insufficient for genome-wide analysis techniques, we examined changes in chromatin organization in cells migrating through 3D collagen matrices with different average pore sizes, achieved by polymerization solutions with different collagen concentrations. We chose collagen concentrations that result in average pore sizes ranging from ≈27 μm^2^ (0.3 mg/ml, low concentration) to ≈4 μm^2^ (1.7 mg/ml, high concentration) in cross section (Denais et al., 2016; Wolf et al., 2013) (Fig. 5A). Experiments were performed either in the absence or presence of a broadband matrix metalloproteinase (MMP) inhibitor. When MMPs are inhibited, cells must migrate through pre-existing pores in the collagen matrix, whose size correlates with the collagen concentration (Wolf et al., 2013).

**Figure 5:**
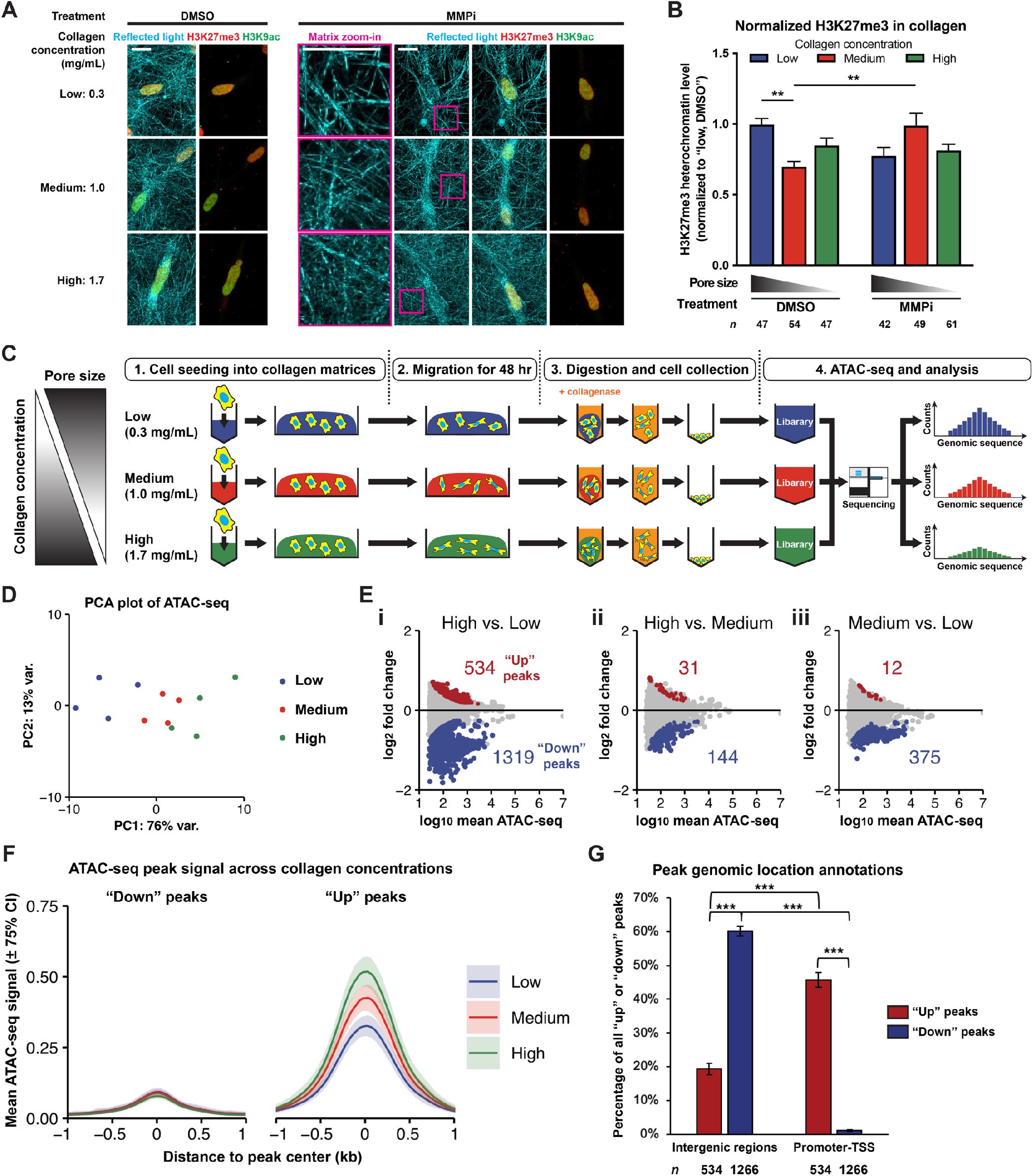
Confined migration alters chromatin accessibility. **(A)** Representative staining of H3K27me3 (red) and H3K9ac (green) in HT1080 cells migrating in low (0.3 mg/mL), medium (1.0 mg/mL) or high (1.7 mg/mL) concentration collagen matrices, under DMSO (vehicle control) or MMPi treatment. Reflected light images of the collagen matrices is shown in cyan. Zoom-in regions of the collagen matrices under MMPi treatment are marked by magenta. Scale bars:15 μm. **(B)** Quantification of normalized H3K27me3 heterochromatin level in HT1080 cells migrating in collagen matrices, under DMSO or MMPi treatment. All values are normalized to DMSO-treated cells in low concentration collagen. \*\**p* < 0.01, two-way ANOVA with Tukey’s multiple comparison test. Data are presented as mean ± SEM, based on *n* cells pooled from three biological replicates. **(C)** The workflow of ATAC-seq experiments. HT1080 single cells were seeded into low, medium and high concentration collagen matrices (step 1), and allowed migration for 48 hours (step 2). Collagen matrices were collected, and collagenase was added for digestion and collection of the cells (step 3). ATAC-seq libraries constructed from all samples were sent for sequencing and the results were analyzed (step 4). **(D)** Principal component analysis (PCA) of all samples. Numbers on PC1 and PC2 axes: percentage of variance (var.) explained by the principal component. **(E)** MA (logged intensity ratio — mean logged intensities) plots of (i) high versus low, (ii) high versus medium, and (iii) medium versus low concentration collagen samples. Y-axis: log_2_-fold changes of the peak signal, with the higher concentration collagen samples divided by the lower concentration collagen samples. X-axis: the mean of log ATAC-seq signal (sequencing reads) across the four biological replicates. Red: up-regulated differentially accessible (DA) peaks (“up” peaks). Purple: down-regulated DA peaks (“down” peaks). Numbers next to the highlighted data points: the numbers of DA peaks. **(F)** Meta-analysis of ATAC-seq peak signal across collagen concentrations, which compiles the mean of ATAC-seq peak signal in all samples of “down” and “up” peaks (from the high versus low concentration collagen samples). Curves and shaded areas represent mean ± 75% confidence interval (CI). Y-axis: ATAC-seq signal. X-axis: distance to peak center in kb. **(G)** Annotations of genomic locations that are intergenic or promoter-TSS within “up” and “down” peaks (from the high versus low concentration collagen samples). \*\*\**p* < 0.001, two-way ANOVA with Tukey’s multiple comparison test. Data are presented as mean ± SEM, based on *n* DA peaks.

In contrast to the experiments using microfluidic devices to control cell confinement, cell migration in high concentration 3D collagen matrices did not increase global heterochromatin levels when compared to cells in low concentration collagen matrices, regardless of MMP inhibitor (MMPi) treatment (Fig. 5A, B; Fig. S7A). Furthermore, the degree of nuclear deformation (nuclear circularity) had no correlation with heterochromatin levels (Fig. S7B), although this could also reflect the fact that nuclear deformations are very dynamic and thus images provide only a static snapshot of the current nuclear morphology, whereas changes in heterochromatin levels are much more persistent (Fig. 2). In collagen matrices made with even higher collagen concentrations (2.4 and 3.1 mg/mL), cells migrating in the absence of MMPi showed a trend toward reduced heterochromatin levels under vehicle control; however, since heterochromatin levels remained essentially unchanged in MMPi treated cells, independent of collagen concentration (Fig. S7C), this trend could not be attributed to differences in confinement, and may instead reflect other effects of increased collagen concentrations in 3D matrices. Surprisingly, increasing collagen concentrations in 2D culture did not affect heterochromatin levels (Fig. S7D). Overall, these results suggest that when cells migrate through confined environments in 3D collagen matrices under MMPi, global heterochromatin levels do not show an increase, similar to previous reports (Golloshi et al., 2020; Wang et al., 2018). At the same time, these findings point to the difficulty in interpreting results from 3D collagen matrices due to the pleiotropic effects of varying collagen concentrations. For example, the discrepancy between the immunofluorescence staining results for global changes of heterochromatin levels obtained in the microfluidic devices versus the 3D collagen matrices could result from effects of increased collagen concentrations on the viscoelastic properties of the matrix or increased collagen-cell surface receptor interactions.

Despite the lack of overt differences in global heterochromatin levels, we reasoned that cells subjected to confined migration may still exhibit local chromatin changes undetected by whole-cell staining levels, as reported by Golloshi et al. (Golloshi et al., 2020). To test this hypothesis, we performed Assay for Transposase-Accessible Chromatin using sequencing (ATAC-seq) analysis, which measures the accessibility of the chromatin genome-wide by transposase-based tagging and fragmentation of the genome (Corces et al., 2017). Experiments were performed in the presence of MMPi treatment to ensure confined pore sizes in the matrices. We designed an ATAC-seq pipeline involving cell collection out of the collagen matrices using collagenase digestion (Fig. 5C), which preserved heterochromatin levels (Fig. S7E). Principal component analysis (PCA) of ATAC-seq data showed clustering of samples according to the collagen concentrations (Fig. 5D), suggesting the progressive effect of increasing collagen concentrations (and decreasing pore size) on local chromatin accessibility. The open and accessible euchromatin is expected to exhibit increased accessibility (i.e., high transposase-tagged fragment reads and therefore high ATAC-seq signal), while compact and inaccessible heterochromatin is expected to show decreased accessibility (i.e., low transposase-tagged fragment reads and therefore low ATAC-seq signal). Differential accessibility analysis of high versus low concentration collagen samples revealed 534 differentially accessible (DA) peaks that were up-regulated in high concentration collagen samples (referred to as “up” peaks), and 1,319 down-regulated DA peaks (“down” peaks) in high concentration collagen samples (Fig. 5E, panel i). The larger number of “down” peaks (purple) compared to “up” peaks (red) indicates an overall trend of reduction in chromatin accessibility, consistent with the global increase in H3K27me3 and H3K9me3 in the microfluidic devices and suggesting that in cells migrating in high concentration collagen matrices still exhibit local and small-scale heterochromatin formation, despite the lack of global increase. Other comparisons, i.e., high vs. medium and medium vs. low collagen concentrations, showed similar but less pronounced patterns, with a larger number of “down” than “up” peaks (Fig. 5E, panels ii-iii). A review of all “down” peaks, aligned based on their peak centers, revealed that the ATAC-seq signal of these peaks remained low at all collagen concentrations, without large visual differences, despite their statistical significance (Fig. 5F, left panel). These data suggest that these peaks correspond to genomic regions that are already silenced at baseline and thus only poorly accessible. In contrast, the ATAC-seq signal of “up” peaks progressively increased with increasing collagen concentration (Fig. 5F, right panel), suggesting that “up” peaks may be regulating genes activated by confined migration. As expected, when we quantified the genomic location annotation of these DA peaks, we found that the majority of “down” peaks were located at intergenic regions, whereas most of the “up” peaks were located at promoter-transcription start sites (TSS) of known genes (Fig. 5G).

### Confined migration decreases intergenic chromatin accessibility near centromeres and telomeres

To further access the potential effects of chromatin accessibility changes on nearby genes, we plotted each peak’s distance to the TSS of its associated genes (i.e., computationally mapped genes and regulatory domains within ±1,000 kb of the peak) using Genomic Regions Enrichment of Annotations Tool (GREAT) (McLean et al., 2010). Most of the “down” peaks were located far (i.e., more than >50 kb, a common long-distance cutoff (Akhtar et al., 2013; Wong, 2016)) from the TSS of the associated genes (Fig. 6A), consistent with their annotation as being located in “intergenic regions” (Fig. 5G). These data suggest that most of the computationally associated genes are unlikely to be regulated by “down” peaks, although in some cases, ATAC-seq signals of the “down” peaks mapped onto the corresponding genes displayed small but progressive decrease in accessibility from low to high collagen concentrations (Fig. 6B; Table S1).

**Figure 6:**
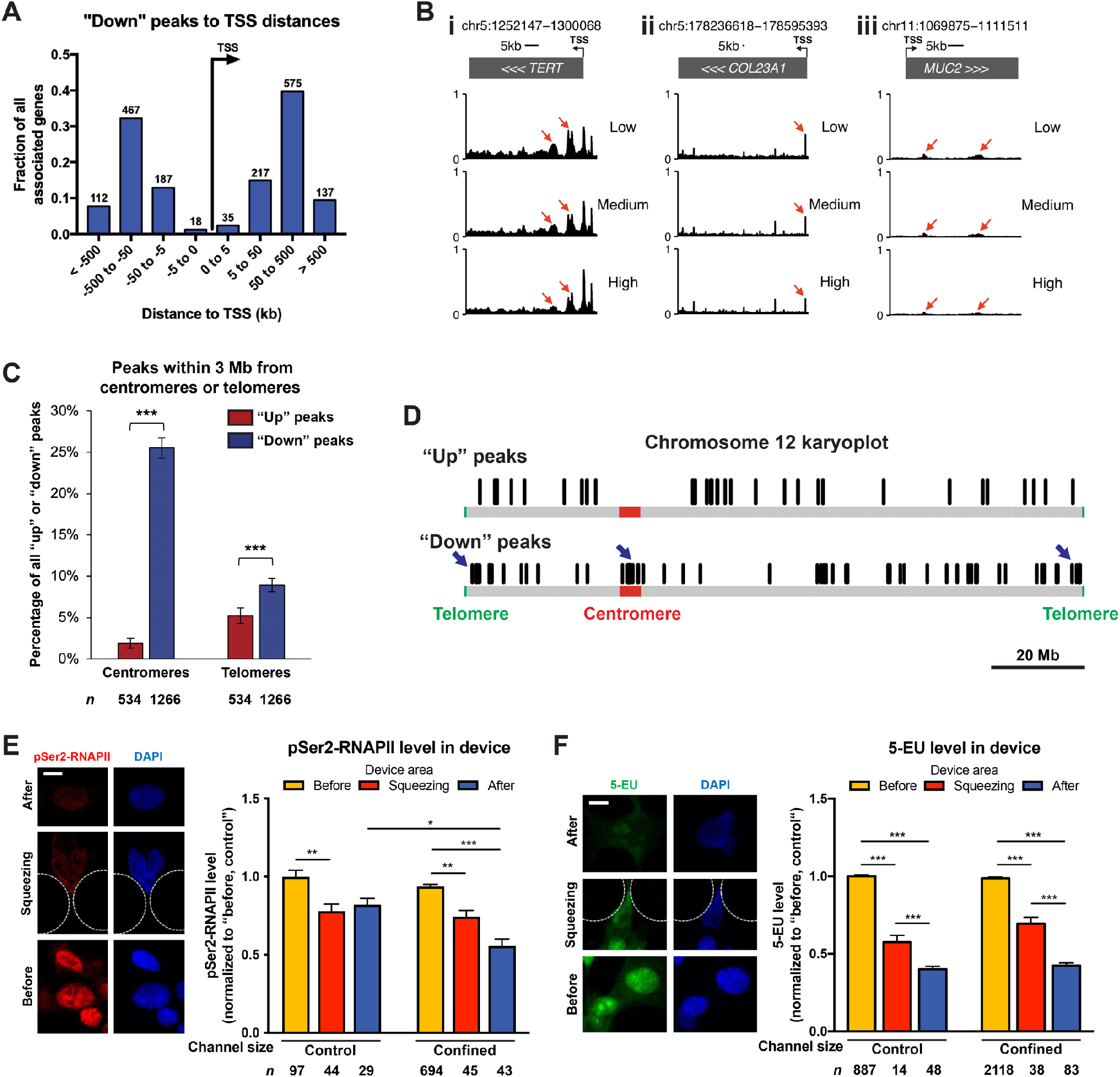
Confined migration decreases intergenic chromatin accessibility near centromeres and telomeres, and global transcription. **(A)** Distribution of distance between each “down” peak and TSS of its associated genes, calculated by GREAT annotated region-gene associations. Y-axis: the fraction of all associated genes. X-axis: binned distances to TSS in kb. The number on top of each bar: the number of peak-associated genes with distances in that bin. **(B)** Genome browser shots of three representative genes associated with “down” peaks, (i) *TERT*, (ii) *COL23A1*, and (iii) *MUC2* ATAC-seq signals in low, medium and high concentration collagen samples. Red arrows: ATAC-seq peaks that are negatively correlated with increasing collagen concentrations. **(C)** The percentage of “up” and “down” peaks located within 3 Mb from centromeres or telomeres. \*\*\**p* < 0.001, two-way ANOVA with Tukey’s multiple comparison test. **(D)** Representative karyoplot of DA peaks on chromosome 12. Black lines: locations of “up” peaks (top) and “down” peaks (bottom). Red areas: centromeres. Green areas: telomeres. Blue arrows: clustering of “down” peaks near centromeres and telomeres. **(E)** Left panel: Representative images of pSer2-RNAPII (red) and DAPI (blue) in HT1080 cells during confined migration. Scale bar: 10 μm. Right panel: Quantification of pSer2-RNAPII intensities in cells migrating through control or confined channels. All values are normalized to control channels “before” cells. \**p* < 0.05, \*\**p* < 0.01, \*\*\**p* < 0.001, two-way ANOVA with Tukey’s multiple comparison test. **(F)** Left panel: Representative images of 5-EU (green) and DAPI (blue) in HT1080 cells during confined migration. Scale bar: 10 μm. Right panel: Quantification of 5-EU intensities in cells migrating through control or confined channels. All values are normalized to control channels “before” cells. \*\*\**p* < 0.001, two-way ANOVA with Tukey’s multiple comparison test. Data are presented as mean ± SEM, based on *n* DA peaks (or *n* cells, listed in each graph) pooled from three biological replicates, except for data in (E), which are based on two biological replicates.

Our findings indicate that most “down” peaks are in genomic regions with low chromatin accessibility, i.e., regions that are likely already silenced by heterochromatin at baseline. We thus hypothesized that heterochromatin spreading from centromeric or telomeric regions plays a role in CMiH formation, since heterochromatin spreading from pre-existing heterochromatin is a common mechanism of heterochromatin formation (Allshire and Madhani, 2018; Wang et al., 2014). When we mapped the distance of each DA peak to the centromere or telomeres of the same chromosome, we found that many “down” peaks, but only few “up” peaks, were located within 3 Mb of centromeres or telomeres (Fig. 6C), well within the < 9 Mb definition of peri-centromeric regions or the < 7 Mb definition of sub-telomeric regions (Levy-Sakin et al., 2019). When mapping peaks to whole chromosomes as karyoplots, “down” peaks were visually clustered within or near the centromeres and telomeres, whereas “up” peaks did not follow any obvious pattern (Fig. 6D; Fig. S8A, B). Moreover, “down” peaks shared among all three different comparisons (high vs. low, high vs. medium, and medium vs. low) of collagen concentration samples (Fig. S8C) had even more peaks within 3 Mb of centromeres or telomeres than “down” peaks in high vs. low comparison only (Fig. S8D), suggesting that CMiH formation consistently clustered around these constitutive heterochromatin sites.

Since centromeres, telomeres, and nearby regions contain an abundance of highly repetitive DNA sequences and transposable elements (TEs) (Slotkin and Martienssen, 2007), which are not detectable by conventional sequencing analysis methods, we used custom-developed algorithms (Kapusta et al., 2013; Lynch et al., 2015) to map TEs to “up” or “down” peaks. As expected, “down” peaks were significantly enriched with TEs, including the cancer-overexpressed alpha satellite, ALR_Alpha (Bersani et al., 2015), and one of the most active TEs in the abundant Alu TE family, AluYj4 (Bennett et al., 2008), (Fig. S8E, panel i). In contrast, “up” peaks were depleted for TEs (Fig. S8E, panel ii). The locations of the enriched TEs matched centromeres or telomeres (Fig. S8F, G), further strengthening the observation of “down” peaks clustering around these sites. Taken together, these data suggest that CMiH is associated with reduced chromatin accessibility near centromeres and telomeres, likely resulting from spreading of constitutive heterochromatin sites, rather than the silencing of new and gene-rich genomic regions.

### Confined migration decreases global transcription and nascent mRNA levels

Since the global increase in heterochromatin marks and the ATAC-seq analysis suggest a predominantly repressive effect of confined migration on gene expression, we investigated the impact of confined migration on RNA polymerase II phosphorylation (pSer2-RNAPII), a marker of active transcription elongation (Bartkowiak and Greenleaf, 2011). Consistent with the chromatin modifications, cells that had migrated through either the control or confined channels had lower pSer2-RNAPII levels than cells in the unconfined “before” area, but the effect was more pronounced in the confined channels (Fig. 6E), indicating that confined migration represses transcription. To directly visualize the relative level of transcription, we pulsed cells with 5-ethynyl uridine (5-EU) to label nascent mRNA transcripts (Jao and Salic, 2008). The results revealed a significant decrease of nuclear 5-EU signal in cells squeezing through the constrictions and after migration through the control and confined channels (Fig. 6F), consistent with the reduced RNAPII phosphorylation and a decrease in transcription. The fact that even migration through the 15× 5 µm^2^ control channels reduced transcription was consistent with our earlier finding that the 5 μm channel height is sufficient for CMiH formation (Fig. 1H, I). The overall repressed transcription in cells migrating through 5-μm tall channels supports the ATAC-seq results of local repressive chromatin accessibility changes induced by confined migration, reflecting the silencing effect of CMiH.

### Confined migration increases chromatin accessibility at genes associated with diverse cellular functions

Given that a high fraction of “up” peaks locations were annotated as promoter-TSS regions (Fig. 5G), we examined their relationship to genes within ±1,000 kb of the peaks. The peak-to-TSS distribution showed that most of the “up” peaks were located within 5 kb of the TSS of associated genes (Fig. 7A). Mapping the ATAC-seq signals of representative “up” peaks onto the corresponding genes revealed a progressive increase in accessibility near the genes’ promoter-TSS from low to high concentration collagen (Fig. 7B; Table S1). However, although statistically highly significant, the log_2_-fold changes (log2FC) of “up” peaks were typically small in magnitude, with the largest log_2_-fold changes only ≈`0.4 (Table S1), suggesting that any associated transcriptional changes may be relatively small and difficult to detect. Accordingly, we did not find any significant difference in the gene expression during migration in 3D collagen matrices or protein levels during confined migration in the microfluidic devices of two representative genes associated with “up” peaks, *HDAC3* and *CBX5/HP1α* (Fig. S10 C-F). However, we cannot rule out the possibility that other “up” peaks-associated genes may show actual transcriptional changes.

**Figure 7:**
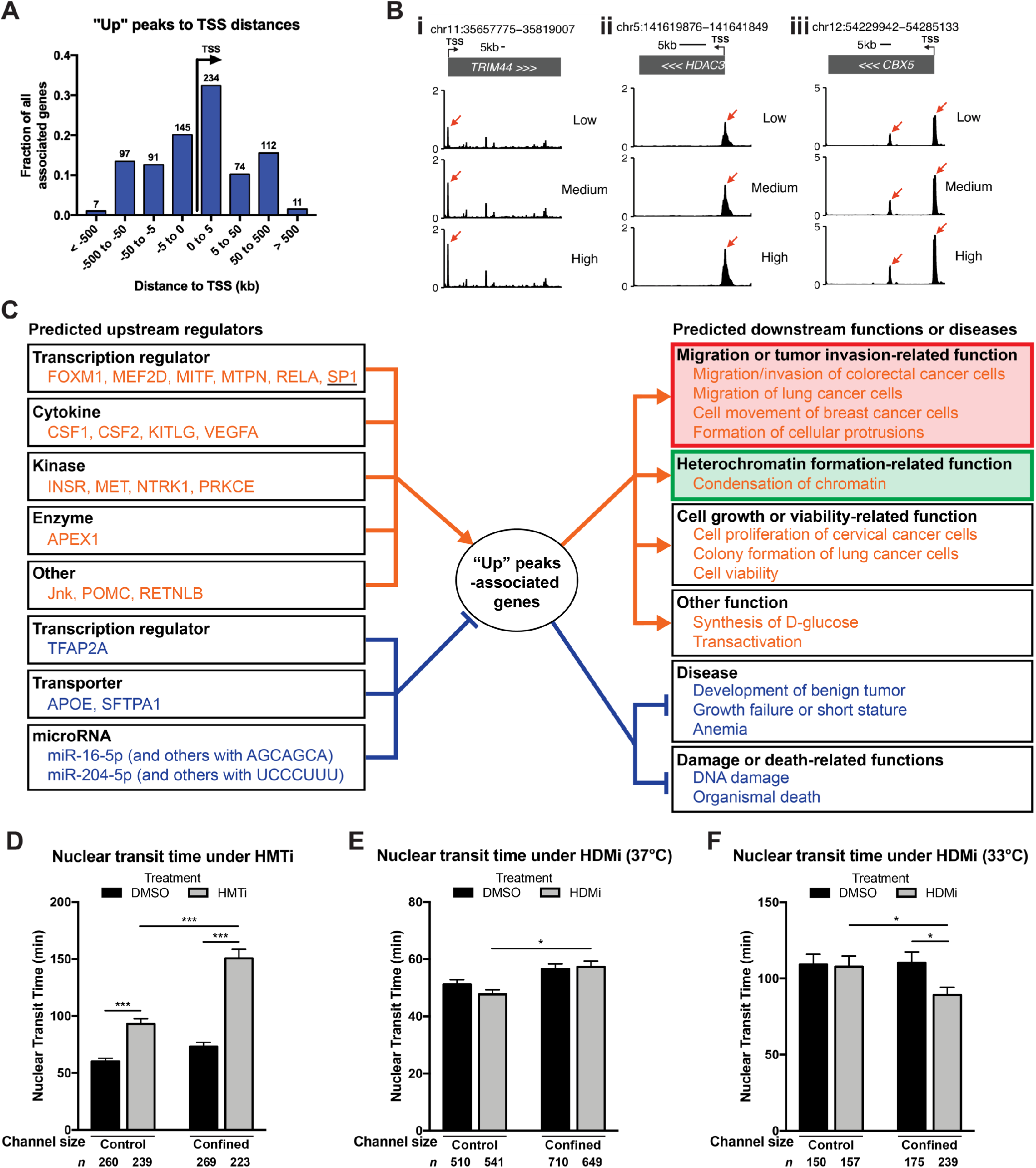
Confined migration increases chromatin accessibility at genes of diverse functions, and confined migration is promoted by heterochromatin formation. **(A)** Distribution of distance between each “up” peak and TSS of its associated genes, calculated by GREAT annotated region-gene associations. Y-axis: the fraction of all associated genes. X-axis: binned distances to TSS in kb. The number on top of each bar: the number of peak-associated genes with distances in that bin. **(B)** Genome browser shots of three representative genes associated with “up” peaks, (i) *TRIM44*, (ii) *HDAC3*, and (iii) *CBX5/HP1α* ATAC-seq signals in low, medium and high concentration collagen samples. Red arrows: ATAC-seq peaks that are positively correlated with increasing collagen concentrations. **(C)** Predicted network of upstream regulators (left) and downstream functions (right) of genes associated with “up” peaks, as calculated by IPA (Ingenuity Pathway Analysis). Orange indicates activation, whereas blue indicates inhibition. Underlined: SP1. Red box: activation of cell migration or tumor invasion-related functions. Green box: activation of heterochromatin formation-related function. For the full network, please refer to Figure S9. **(D)** Quantification of HT1080 cells nuclear transit time through control or confined channels, under DMSO (vehicle) or HMTi treatment. \*\*\**p* < 0.001, Kruskal-Wallis H test with Dunn’s multiple comparison test. Please see also Video S4-S7. **(E)** Quantification of HT1080 cells nuclear transit time through control or confined channels, under DMSO (vehicle) or HDMi treatment. \**p* < 0.05, Kruskal-Wallis H test with Dunn’s multiple comparison test. **(F)** Quantification of HT1080 cells nuclear transit time through control or confined channels, under DMSO (vehicle) or HDMi treatment under mild hypothermia stress (at 33 °C). \**p* < 0.05, Kruskal-Wallis H test with Dunn’s multiple comparison test. Please see also Video S8, S9. Data are presented as mean ± SEM, based on *n* cells (listed in each graph) pooled from at least three biological replicates.

To explore the potential functional consequences of genes associated with “up” peaks, we performed Gene Ontology (GO) Biological Process enrichment analysis of these genes (Table 1), revealing an enrichment in chromatin/gene silencing and other chromatin structure-related functions (denoted with * in Table 1), DNA damage checkpoint (Table S3), and cell cycle checkpoint (Table S4). Moreover, GO Cellular Component enrichment analysis revealed enrichment of DNA packaging complex, nucleosome, and chromosome (telomeric region) associated genes (Table S5). Interestingly, many of the annotated genes encode histones proteins, including genes from all four major histone families of H2A, H2B, H3 and H4, as well as several genes critical for heterochromatin formation and maintenance, such as histone acetyltransferase 1 (*HAT1*), *HDAC3*, and *CBX5* (which encodes HP1α) (denoted with * in Table S2).

**Table 1:**
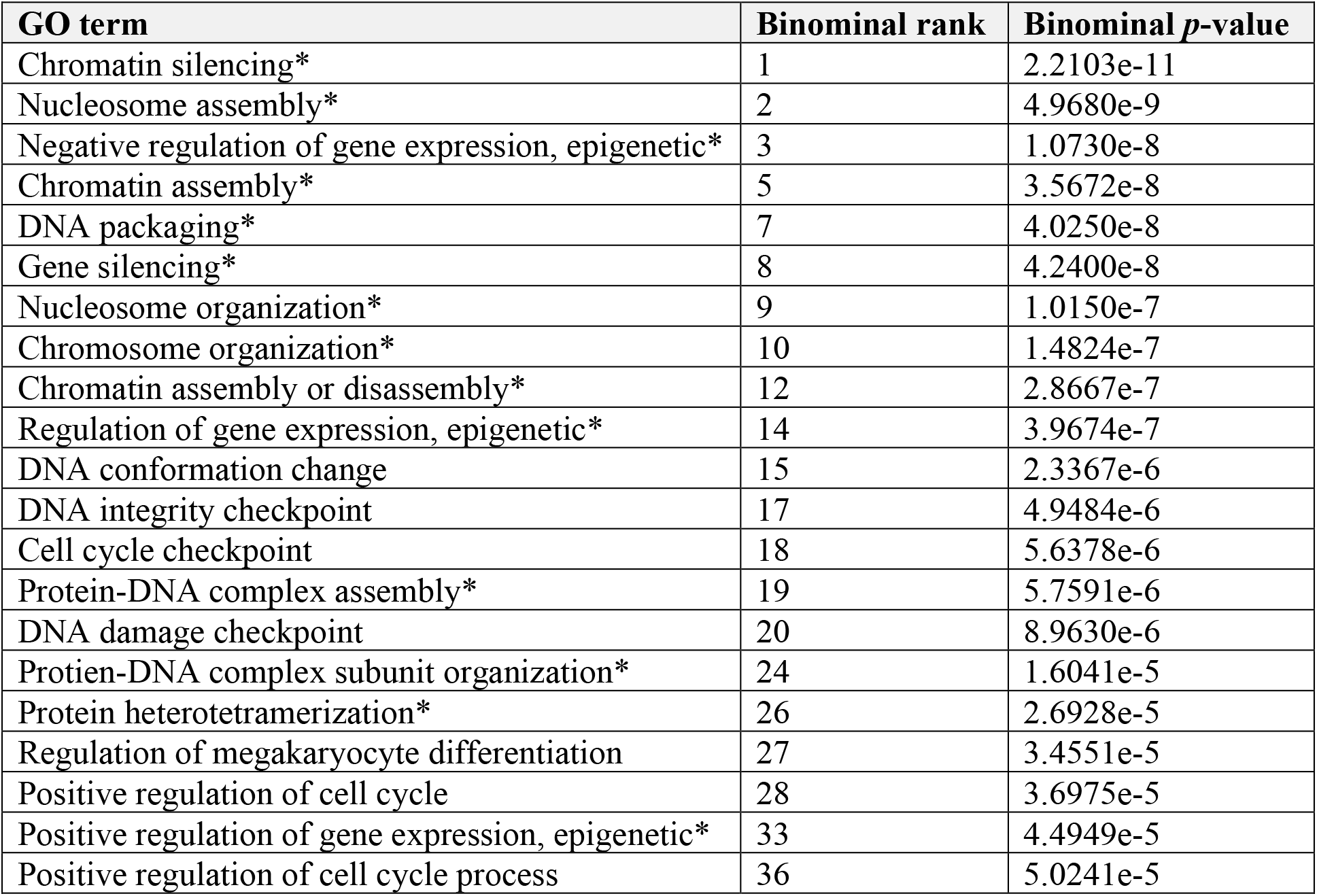
Gene Ontology (GO) Biological Process terms enrichment of genes associated with up-regulated DA peaks in high concentration collagen samples. *Chromatin structure or gene silencing-related GO terms.

| GO term | Binominal rank | Binominal *p*-value |
| --- | --- | --- |
| Chromatin silencing* | 1 | 2.2103e-11 |
| Nucleosome assembly* | 2 | 4.9680e-9 |
| Negative regulation of gene expression, epigenetic* | 3 | 1.0730e-8 |
| Chromatin assembly* | 5 | 3.5672e-8 |
| DNA packaging* | 7 | 4.0250e-8 |
| Gene silencing* | 8 | 4.2400e-8 |
| Nucleosome organization* | 9 | 1.0150e-7 |
| Chromosome organization* | 10 | 1.4824e-7 |
| Chromatin assembly or disassembly* | 12 | 2.8667e-7 |
| Regulation of gene expression, epigenetic* | 14 | 3.9674e-7 |
| DNA conformation change | 15 | 2.3367e-6 |
| DNA integrity checkpoint | 17 | 4.9484e-6 |
| Cell cycle checkpoint | 18 | 5.6378e-6 |
| Protein-DNA complex assembly* | 19 | 5.7591e-6 |
| DNA damage checkpoint | 20 | 8.9630e-6 |
| Protein-DNA complex subunit organization* | 24 | 1.6041e-5 |
| Protein heterotetramerization* | 26 | 2.6928e-5 |
| Regulation of megakaryocyte differentiation | 27 | 3.4551e-5 |
| Positive regulation of cell cycle | 28 | 3.6975e-5 |
| Positive regulation of gene expression, epigenetic* | 33 | 4.4949e-5 |
| Positive regulation of cell cycle process | 36 | 5.0241e-5 |

Canonical pathway analysis of the genes associated with “up” peaks (i.e. those whose TSS was nearest to the peak) using Ingenuity Pathway Analysis (IPA) (Krämer et al., 2014) identified a diverse list of significant pathways, including pathways related to cell migration, tumor invasion, histone methylation, and heterochromatin formation (Table 2). Moreover, several of the pathways, such as TGF-β signaling, Pyridoxal 5’-phosphate Salvage Pathway and ERK5 Signaling, had previously been linked to migration-induced H3K27me3 heterochromatin in scratch wound assays (Segal et al., 2018). Predicted regulatory networks of “up” peak-associated genes generated using IPA identified cell migration or tumor invasion, heterochromatin formation, and cell growth among the activated downstream functions, whereas DNA damage was among the inhibited downstream functions (Fig. 7C; Fig. S9). Moreover, genes associated with “up” peaks that were shared among different comparisons (high vs. low, high vs. medium, and medium vs. low) of collagen concentration samples (Fig. S10A) exhibited the same predicted downstream functions (Fig. S10B), suggesting that such gene functions are important for confined migration. Overall, the presence of numerous chromatin structure-related functions in the GO term and pathway analysis indicates that changes in chromatin accessibility caused by confined migration may drive further chromatin remodeling steps involved in CMiH. Other cellular functions identified in the analysis are consistent with previous findings that cell migration activates tumor invasion and cell proliferation pathways(Segal et al., 2018), and that migration through confined spaces can induce DNA damage pathways (Denais et al., 2016; Irianto et al., 2017, 2016; Shah et al., 2020) and alter cell cycle progression (Moriarty and Stroka, 2018; Xia et al., 2019).

**Table 2:** Canonical pathways of genes associated with up-regulated DA peaks in high concentration collagen samples. Only pathways with -log (*p*-value) > 1.30 (*p*-value < 0.05) and |Z-score| (if known) ≥ 1 are shown. N/A, not available. †Cell migration or tumor invasion-related pathways. *Histone methylation or heterochromatin formation-related pathways.

| Ingenuity Canonical Pathways | Rank | $-\log(p\text{-value})$ | Z-score |
| --- | --- | --- | --- |
| NRF2-mediated Oxidative Stress Response† | 1 | 3.94 | 2.333 |
| Antioxidant Action of Vitamin C | 2 | 3.22 | -2.449 |
| Spliceosomal Cycle | 3 | 2.86 | 2.236 |
| Kinetochore Metaphase Signaling Pathway | 4 | 2.74 | 1.134 |
| Protein Ubiquitination Pathway | 5 | 2.54 | N/A |
| Superpathway of Methionine Degradation* | 6 | 2.44 | 2 |
| mTOR Signaling† | 7 | 2.44 | 1.633 |
| p38 MAPK Signaling† | 8 | 2.36 | 1.342 |
| Estrogen Receptor Signaling† | 9 | 2.32 | 2.887 |
| Phospholipases | 10 | 2.28 | 2.236 |
| Thioredoxin Pathway | 11 | 2.23 | N/A |
| Hypoxia Signaling in the Cardiovascular System | 12 | 2.05 | N/A |
| Cell Cycle: G2/M DNA Damage Checkpoint Regulation | 13 | 2 | -2 |
| Toll-like Receptor Signaling | 14 | 2 | N/A |
| CD27 Signaling in Lymphocytes | 15 | 1.88 | 2 |
| Endothelin-1 Signaling† | 16 | 1.78 | 2.828 |
| Role of PKR in Interferon Induction and Antiviral Response | 17 | 1.78 | 1.342 |
| Glycogen Degradation II | 18 | 1.76 | N/A |
| IL-1 Signaling† | 19 | 1.67 | 1.342 |
| DNA Methylation and Transcriptional Repression Signaling* | 20 | 1.66 | N/A |
| Glycogen Degradation III | 21 | 1.63 | N/A |
| TGF-β Signaling† | 22 | 1.6 | 1.342 |
| Insulin Secretion Signaling Pathway | 23 | 1.58 | 3 |
| Choline Biosynthesis III | 24 | 1.57 | N/A |
| Mitotic Roles of Polo-Like Kinase | 25 | 1.57 | N/A |
| Pyridoxal 5'-phosphate Salvage Pathway† | 26 | 1.57 | N/A |
| UVA-Induced MAPK Signaling | 27 | 1.56 | N/A |
| Remodeling of Epithelial Adherens Junctions† | 28 | 1.53 | N/A |
| GNRH Signaling | 29 | 1.51 | 2.646 |
| April Mediated Signaling | 30 | 1.5 | N/A |
| IL-10 Signaling† | 31 | 1.49 | N/A |
| NER Pathway | 32 | 1.48 | -1.342 |
| B Cell Activating Factor Signaling | 33 | 1.48 | N/A |
| IGF-1 Signaling† | 34 | 1.47 | N/A |
| ERK5 Signaling† | 35 | 1.45 | 2 |
| Endocannabinoid Cancer Inhibition Pathway | 36 | 1.42 | 1.633 |
| Pancreatic Adenocarcinoma Signaling† | 37 | 1.39 | N/A |
| iNOS Signaling† | 38 | 1.37 | N/A |
| PFKFB4 Signaling Pathway† | 39 | 1.35 | N/A |
| Endocannabinoid Developing Neuron Pathway | 40 | 1.31 | 1 |
| Renal Cell Carcinoma Signaling† | 41 | 1.31 | N/A |
| Chemokine Signaling† | 42 | 1.31 | 2 |

To identify transcription factors (TFs) that may regulate “up” peaks in response to confined migration, we performed motif enrichment analysis on “up” peak sequences and matched the enriched DNA motifs to known TF binding motifs (Fig. S10G). The identified TF included SP1 and SMAD3 (underlined in Fig. S10G), which form a complex under TGF-β signaling to regulate tumor progression and invasion (Jungert et al., 2006; Lang et al., 2020). Intriguingly, SP1 was also a predicted upstream regulator (underlined in Fig. 7C), whereas *SMAD3* gene was identified as part of the chromatin GO terms (denoted with † in Table S2) and one of the genes associated with the shared “up” peaks (highlighted in Fig. S10B). These findings suggests that both SP1 and SMAD3 could serve potential roles as upstream TFs that regulate accessibility and downstream functions in genes associated with “up” peaks.

### Confined migration regulates co-accessibility of enhancer-promoter interactions

One central mechanism by which chromatin accessibility changes can regulate gene expression is through enhancer-promoter interactions (EPIs) that exhibit co-accessibility, i.e., having the same direction of accessibility changes (Klemm et al., 2019; Pliner et al., 2018). To identify potential EPIs in “up” and “down” peaks, we assigned peaks that were within 1 kb upstream of the nearest gene’s TSS to be “promoter-associated”, and all other peaks as “non-promoter-associated”. As expected, we identified many more promoter-associated peaks in “up” peaks than in “down” peaks (Fig. S11A). We then mapped all peaks to the DNase I hypersensitive regions in the ENCODE database (Rosenbloom et al., 2013), as DNase I hypersensitivity validates chromatin regulatory elements, including promoters and enhancers (Tsompana and Buck, 2014). “Up” peaks overlapped much more with the DNase I data, suggesting that “up” peaks contained more putative promoters and enhancers than “down” peaks (Fig. S11B). Based on the validated peaks, we found that EPI distance (promoter to non-promoter, peak-to-peak distance) was significantly longer for “up” peaks, compared to “down” peaks (Fig. S11C). This finding suggests that long-range (i.e., more than a few hundreds of kb) enhancer-mediated regulation plays a role in “up” peaks-associated genes, which is a sign of epigenetic dysregulation and compromised chromatin domains in cancer (Schoenfelder and Fraser, 2019). We identified 41 and 7 EPIs in “up” peaks and “down” peaks, respectively, based on the conventional 500 kb cutoff (van Arensbergen et al., 2014) (Fig. S11D). Representative EPIs in “up” peaks mapped to the enhancer and promoter database annotations (Fig. S11F), supporting the validity of our EPI identification. Overall, these identified EPIs suggest that confined migration-induced chromatin accessibility changes may influence gene expression via enhancer-mediated regulation.

### Heterochromatin formation promotes confined migration

The significant extent of CMiH formation, the associated local chromatin accessibility changes, and the predicted potential downstream functions prompted us to investigate effects of CMiH on migration. Previous studies showed that reducing heterochromatin levels by treatment with pan-HDACi (TSA) or HMTi (5’-deoxy-5’-methylthioadenosine) impairs migration through 3D confined environments (Fu et al., 2012; Krause et al., 2019). Since reduced nuclear stiffness via lamin depletion facilitates nuclear transit through confined spaces (Davidson et al., 2014; Lautscham et al., 2015), it is puzzling that the reduced nuclear stiffness due to lowered heterochromatin level would have an opposite effect on confined migration (Krause et al., 2019). Therefore, we decided to test if CMiH offered an advantage for cell migration through confined channels. HMTi (DZNep) treatment resulted in increased nuclear transit time (i.e., decreased migration speed) through constrictions in both 15×5 μm^2^ control and ≤2×5 μm^2^ confined channels compared to vehicle controls, but the effect was less pronounced in control channels (Fig. 7D; Video S4-S7). The inhibitory effect of HMTi treatment on migration speed was even less in cells migrating through 10-μm tall (15×10 μm^2^) channels (Fig. S12A), which do not confine the nucleus. The progressively increasing effect of HMTi treatment on nuclear transit times when comparing 10-μm tall channels, 5-μm tall control channels, and the confined channels closely mirrors our earlier findings that 5-μm tall control channels and confined channels result in increasing levels of heterochromatin compared to the 10-μm tall channels (Fig. 1D, F, I), demonstrating that CMiH is particularly beneficial for migration in increasingly confined conditions. This effect of HMTi on confined migration speed was consistent with previous studies (Fu et al., 2012; Krause et al., 2019), demonstrating the importance of CMiH for efficient nuclear transit through tight spaces. On the other hand, increasing heterochromatin during confined migration via HDMi (JIB-04) treatment did not affect migration speed (Fig. 7E). However, under conditions that slow down migration, such as mild hypothermia (33°C), which inhibits cancer cell proliferation and migration (Fulbert et al., 2019; Kalamida et al., 2015), HDMi reduced nuclear transit time through constrictions in confined channels compared to vehicle controls (Fig. 7F; Video S8, S9). These data suggest that additional heterochromatin formation does not speed up confined migration in general, but that it can promote confined migration under adverse conditions. Overall, our data show that CMiH is crucial for efficient confined migration, and additional heterochromatin can promote confined migration under environmental conditions adverse for cell migration.

## Discussion

In this study, we demonstrated that CMiH is a prominent feature across cancer and non-cancerous cell lines. Confined migration in microfluidic devices led to increases in H3K27me3 and H3K9me3 levels, both globally and in nuclear blebs, but H3K27me3 increased more substantially, resonating with previous findings of its importance in cancer cell migration (Liu et al., 2018, p. 2; Segal et al., 2018, p. 27). Vertical confinement to 5 µm height was sufficient for CMiH formation, consistent with previous studies on the nuclear mechanosensing threshold being around 5 to 7 μm (Lomakin et al., 2020; Venturini et al., 2020), and CMiH increased further with increasing confinement and with migration through successive constrictions. An alternative explanation of the observed chromatin changes is that they are permissive for confined migration, and thus cells with these features are passively selected for during confined migration. Although time-lapse microscopy revealed formation of persistent GFP-HP1α enrichments in cells during and after nuclear transit through tight constrictions, indicating an active heterochromatin formation process, and the vast majority of cells that had passed through the constrictions had higher normalized H3K27me3 levels than observed in any cell in the “before” population (Fig. S1C), we cannot rule out the possibility that the observed increase in heterochromatin is a combination of both active induction and passive selection, as suggested by Golloshi et al. regarding the chromatin changes resulted from repeated transwell migration (Golloshi et al., 2020). Taken together, these findings indicate that CMiH is distinct from heterochromatin associated with general migration initiation, which occurs prior to the onset of migration in 2D and 3D environments (Gerlitz and Bustin, 2010; Segal et al., 2018).

Although we detected significant increases in global heterochromatin levels in cells migrating through confined microfluidic channels, we did not observe the same global changes in cells migrating through 3D collagen matrices with decreasing average pore sizes. The discrepancy between the microfluidic devices and collagen matrices is likely due to the more complex factors contributing to chromatin changes in collagen matrices, such as matrix stiffness (Zhao et al., 2021) and integrin binding to matrix ligands (Carley et al., 2021), whereas the microfluidic devices allow control of cell confinement without affecting the substrate stiffness or ligand density. This precise control of confinement alone makes microfluidic devices an ideal model for studying confinement-induced biological responses.

Intriguingly, CMiH persisted for at least 5 days and was inheritable after mitosis. One possible consequence of these increased heterochromatin levels is replication stress and subsequent DNA damage (Jiang et al., 2009; Kurashima et al., 2020; Lambert and Carr, 2013), which could reduce cell viability. In support of this hypothesis, DNA damage checkpoint and cell cycle checkpoint were among the enriched GO terms associated with increased chromatin accessibility during confined migration. Notably, migration through the confined channels induced around 20% of cell death during a 5-day observation period, compared to 7% cell death for cells migrating through the control channels (Fig. S4D), suggesting that persistent high heterochromatin levels may lead to decreased cell viability, although other factors associated with confined migration could further contribute to the increased cell death. The larger increase in cell death compared to what was reported in a previous study using similar migration devices (Denais et al., 2016) may be due to the shorter observation period (<16 hours) compared to the multi-day time period analyzed here. A recent study found that nuclear deformation during confined migration is sufficient to cause replication stress and DNA damage (Shah et al., 2020), which could then result in heterochromatin formation (Nikolov and Taddei, 2016). However, the study showed that confined migration-induced DNA damage in HT1080 is mostly due to nuclear envelope ruptures (Shah et al., 2020). Here, we showed that CMiH formation is independent of nuclear envelope ruptures (Fig. 4C, D, S6E, F). Thus, it would be difficult to explain the heterochromatin formation purely as a response to DNA damage. Moreover, the study showed that migration through the control channel induces very little DNA damage compared to migration through the confined channels (Shah et al., 2020), while we showed that migration through the control channel still induces a significant amount of heterochromatin formation. In summary, DNA damage response may be involved, but cannot be the major cause of the observed CMiH formation and transcriptional downregulation.

We identified that CMiH is dependent on a combination of histone modifying enzymes, such as HDAC3, and stretch-sensitive ion channels. However, it is unclear if redundant mechanosensing pathways exist for CMiH regulation, such as through the deformation of nuclear-ER membrane and the subsequent calcium ion influx (Lomakin et al., 2020; Venturini et al., 2020), or through force transmission via cytoskeletons and the linker of nucleoskeleton and cytoskeleton (LINC) complexes (Chang et al., 2015; Crisp et al., 2006).

Our ATAC-seq studies suggest that the increase of heterochromatin predominantly corresponds to spreading of existing heterochromatin marks in intergenic or transcriptionally repressed regions. The functional consequences of these modifications could include both changes in the mechanical properties of the nucleus such as nuclear compaction and stiffening (Krause et al., 2019; Stephens et al., 2018), but also transcriptional activity due to decreased enhancer accessibility (Klemm et al., 2019; Pliner et al., 2018). On the other hand, we also identified numerous genes with increased accessibility of their promoter-TSS regions. These genes, which frequently had GO terms and pathways associated with chromatin modifications and tumor cell invasion, could promote both the formation of CMiH and downstream effects of confined migration on other cellular functions. Furthermore, co-accessibility analysis of the ATAC-seq data demonstrated that confined migration may regulate multiple EPIs that leads to gene expression changes, and long-range EPIs may play a role in the regulation of genes with increased accessibility (i.e., genes associated with “up” peaks). However, since there were far less “up” peaks than “down” peaks, the distances between “up” peaks were overall longer (Fig. S11E), potentially resulting in a bias toward overall longer EPI distances in “up” peaks, compared to “down” peaks.

Importantly, we found that CMiH promotes confined migration, as preventing CMiH with HMTi treatment significantly increased transit time, and this effect became increasingly more pronounced with increasing confinement. Possible mechanisms for the pro-migration effect of CMiH include changes in gene expression, nuclear biophysical properties, or a combination of both. For example, CMiH may repress unwanted transcriptional activities, and in turn promote confined migration-related gene expression, similar to the role of H3K27me3 in 2D migration initiation (Segal et al., 2018). On the other hand, CMiH may alter nuclear stiffness and nuclear size (Krause et al., 2019; Stephens et al., 2018), which were shown to affect nuclear transit speed through confined spaces (Krause et al., 2019; Lautscham et al., 2015). Intriguingly, nuclear size was increased under HMTi treatment (Fig. S12B) and decreased under HDMi treatment (Fig. S12C). The exact mechanism by which CMiH promotes confined migration, however, will require further studies beyond the scope of this work. We will further discuss this in the “limitations of the study” section.

In conclusion, we revealed the phenomenon of heterochromatin formation in the context of confined 3D migration, while providing insights on its impact on cell migration and the molecular mechanisms behind it. We have already demonstrated the presence of CMiH in tumor cells and fibroblasts but hypothesize that CMiH also occurs in other cell types, making it relevant for immune responses, wound healing, and during development. Furthermore, the recognition of CMiH in confined migration may provide motivation for future research to target this mechanism to reduce or prevent cancer metastasis. Moreover, our findings could potentially connect the role of chromatin to the role of the nucleus as a cellular mechanosensory (Lomakin et al., 2020; Venturini et al., 2020), and help pave the way for future studies on how chromatin modifications could affect nuclear and cellular responses to the cell’s physical microenvironment.

### Limitations of the study

Since the majority of heterochromatin readouts in this study were based on immunofluorescence labeling at fixed time points, one limitation is the lack of real-time data tracking heterochromatin levels in single cells throughout the migration process with quantifiable reporters such as fluorescence resonance energy transfer (FRET) biosensor (Peng et al., 2018) or mintbodies (Sato et al., 2013) that reports chromatin modification changes. However, such single-cell fluorescence-based methods are very sensitive to the intracellular distribution of the reporter, changes in pH, imaging settings, and phototoxicity/photobleaching during repeated imaging, making them challenging to work with for long-term time-lapse microscopy studies of migrating cells. Supporting assays and analysis that can directly quantify the compaction of chromatin, for example, high magnification imaging of DAPI density and transmission electron microscopy (TEM), would also validate the heterochromatin readout, but are also limited to fixed cells.

The discrepancy between the changes in global heterochromatin levels observed in the microfluidic device versus in 3D collagen matrices is another limitation of this study. As discussed above, multiple factors likely contribute to these conflicting results, and future studies should address whether other approaches to modulate confinement in 3D matrices, such as using different polymerization temperatures, cross-linkers, or engineered hydrogels with customizable specific material properties, can resolve the differences observed here by reducing confounding factors. Although we performed single cell analysis combining immunofluorescence labeling for heterochromatin with reflective light microscopy to confirm the local confinement of individual cells in 3D collagen matrices, we nonetheless acknowledge the difficulties stemming from the heterogeneous nature of collagen matrices, which are expected to particularly affect population measurements, such as analysis used in the ATAC-seq and qPCR experiments. Moreover, the standard normalization methods utilized in our ATAC-seq pipeline may mask the potential global changes of accessibility (Chen et al., 2016; Reske et al., 2020). An external spike-in control (Stewart-Morgan et al., 2019) will enable unbiased detection of global scale changes. The inclusion of a spike-in control is strongly encouraged for follow-up sequencing studies to avoid similar analysis bias. Another limitation of the current ATAC-seq analysis is that we were unable to confirm changes in gene expression or protein levels in the two representative genes we examined. A broader, genome-wide analysis would provide further insights into how confined migration could affect gene expression, but would require collection of larger numbers of cells than currently achievable using our microfluidic devices.

Although our study suggests that confined migration changes chromatin modifications, which may further facilitate cell migration, the exact mechanisms responsible for CMiH and its effect on cell migration remain to be elucidated. Most likely, more than a single mechanism or pathway contribute to the increase in heterochromatin and the associated improvement in confined migration. Aside from potential transcriptional changes (i.e., the predicted regulatory network of ATAC-seq data) and nuclear size that may be affected by histone modifying enzyme inhibitors, we cannot rule out the possibility that other cellular properties, e.g., cell adhesion, traction forces, and cytoskeleton network, may also be affected during the experiments, altering cell migration ability independent of chromatin and nuclear changes.

## STAR Methods

### Resource Availability

#### Lead Contact

Further information and requests for resources and reagents should be directed to and will be fulfilled by the lead contact, Jan Lammerding.

#### Materials Availability

The plasmids and cell lines generated in this study are available from the lead contact on reasonable request.

#### Data and Code Availability

ATAC-seq data that supports the findings of this study have been deposited in Gene Expression Omnibus (GEO) repository, with the accession number GSE181247. The other datasets generated during and/or analyzed during the current study are available from the lead contact on reasonable request.

The ImageJ Macro scripts used to analyze confocal immunofluorescence staining images and to generate normalized heterochromatin images can be accessed at https://github.com/chiehrenhsia/ConfinedMigration. The R scripts used to perform ATAC-seq analyses including the exact parameters can be accessed at https://github.com/jaj256/ConfinedATAC.

### Experimental Model and Subject Details

#### Cell lines and cell culture

The fibrosarcoma cell line HT1080 (ACC315) was purchased from DSMZ in Braunschweig, Germany. The breast adenocarcinoma cell line MDA-MB-231 (ATCC HTB-26) was purchased from American Type Culture Collection (ATCC). The SV40-immortalized human fibroblasts were purchased from the Coriell Institute for Medical Research. All cell lines were cultured in Dulbecco’s Modified Eagle Medium (DMEM, Gibco) supplemented with 10% (v/v) fetal bovine serum (FBS, Seradigm) and 1% (v/v) penicillin and streptomycin (Pen-Strep, Gibco) (i.e., complete DMEM), in the incubator under humidified conditions at 37°C and 5% CO_2_.

#### Generation of fluorescently labeled cell lines

HT1080 cells and MDA-MB-231 cells were stably or transiently modified with either of the following constructs: lentiviral construct GFP-HP1α (pCDH-CMV-EGFP-HP1α-EF1-puro) for imaging of heterochromatin formation; retroviral construct NLS (nuclear localization sequence)-GFP and H2B-tdTomato (pQCXIP-NLS-copGFP-P2A-H2B-tdTomato-IRES-puro) for measuring of nuclear transit time (Elacqua et al., 2018); GFP-HDAC3 (pCMV-EGFP-HDAC3, a gift from Nikhil Jain) for imaging of HDAC3 translocation; lentiviral construct cGAS-mCherry (pCDH-CMV-cGAS^E225A/D227A^-mCherry2-EF1-blastiS) for imaging of NE rupture (Denais et al., 2016).

### Method Details

#### Viral modification

Pseudo-viral particles were produced as described previously (Denais et al., 2016). In brief, 293-TN cells (System Biosciences) were co-transfected with lentiviral packaging plasmid and envelope plasmid (psPAX2 and pMD2.G, gifts from Didier Trono) using PureFection (SBI), following the manufacturer’s protocol. Lentivirus-containing supernatants were collected at 48 hours and 72 hours post-transfection and filtered. Cells were seeded into 6-well plates to reach 50-60% confluency on the day of infection and were transduced 2-3 consecutive days with the viral supernatant in the presence of 8 μg/mL polybrene (Sigma). The viral supernatant was replaced with fresh culture medium, and cells were cultured for 24 hours before selection with 1 μg/mL of puromycin (InvivoGen) or 6ug/mL of blasticidin S (InvivoGen) for 7 days.

#### siRNA-mediated depletion

siRNAs (small interfering RNAs) used were as followed: human *LMNA* (ON-TARGET plus SMARTpool, Dharmachon Horizon, L-004978-00), human *EMD* (Ambion Silencer Select, ID: s4646), human *HDAC3* (ON-TARGET plus SMARTpool, Dharmachon Horizon, L-003496-00), human *CK2α* (Santa Cruz, sc-29918), and non-target (NT) negative control (ON-TARGETplus non-targeting pool, Dharmachon, D-001810-10). Cells were seeded into 12-well plates at a density of 40,000 cells per well the day prior to treatment, and transfected with siRNA using Lipofectamine RNAiMAX (Invitrogen) following the manufacturer’s protocol at a final concentration of 20 nM. Cells were transfected again with fresh siRNA at 24 hours. Cells were trypsinized for migration experiments at 48 hours after transfection. For *HDAC3* knockdown, 40 nM of siRNA was used, and cells were trypsinized for migration experiments at 72 hours after transfection, including 24 hours of recovery in culture media without transfection reagents. Cells treated with the same condition and duration were harvested for validation of successful protein depletion by Western blot analysis and immunofluorescence (IF) staining.

#### Pharmacological treatments

For collagen matrix migration experiments, cells were treated with a board MMP inhibitor, GM6001 (Millipore, 20 μM), to inhibit their matrix degradation ability. For microfluidic migration device experiments, cells were treated with inhibitors at the time of seeding into the migration device and throughout the experiments. Fresh culture media containing inhibitors were changed every 24 hours. Cells were treated with either 5 μM of 3-Deazaneplanocin A (DZNep, Cayman Chemical) for broad-band inhibition of histone methyltransferases (with the exception of 10 μM for HT1080 cells staining experiments); 250 nM of Trichostatin A (TSA, Sigma) for broad-band inhibition of HDACs (with the exception of 125 nM for MDA-MB-231 cells staining experiments); 10 μM of RGFP966 (Selleckchem) for inhibition of HDAC3; 1 μM of JIB-04 (Selleckchem) for inhibition of Jumonji histone demethylases; 10 μM of GdCl_3_ (Santa Cruz) for inhibition of mechanosensitive ions channels. Cells treated with the same condition and duration were harvested for validation of successful enzymatic activity inhibition by Western blot analysis and IF staining. Inhibitor stocks were dissolved in dimethyl sulfoxide (DMSO) (Sigma) (with the exception of GdCl_3_ in water) before diluting into cell culture medium.

#### Single cell collagen matrix migration assays

To create glass-bottom wells for collagen matrices, blocks of polydimethylsiloxane (PDMS) were punched with a 10 mm biopsy punch and covalently bonded onto glass coverslips (VWR) after plasma treatment of 5 minutes. The preparation and making of collagen gel matrices was done as described previously (Cross et al., 2010; Denais et al., 2016). In short, individual wells on glass coverslips were treated with 1% polyethylenimine (PEI, Sigma), followed by 0.1% glutaraldehyde (Sigma) treatment for consistent bonding of collagen matrices, and washed with PBS. To generate single cell-containing collagen matrices at specified collagen concentrations (0.3, 1.0, and 1.7 mg/mL), acidic solution of rat tail type-I collagen (Corning) was supplemented with complete DMEM and NaOH to reach a neutral pH of 7.4, and then mixed with cell suspension to reach a final density of 100,000 cells per mL. Collagen matrices were allowed to polymerize in the incubator for 30 min before adding complete DMEM with either DMSO or a board MMP inhibitor, GM6001 (Millipore, 20 μM), to submerge the matrices. Cells were allowed to migrate in the matrices for 48 hours before fixation and IF staining.

#### Fabrication and use of microfluidic migration devices

The microfluidic migration devices were designed and fabricated as described previously (Davidson et al., 2015; Denais et al., 2016). In brief, PDMS replicas of the migration device molds were made from Sylgard 184 following the manufacturer’s protocol (1:10) and baking at 65°C for 2 hours. Once the PDMS is demolded and cut into individual blocks of devices, biopsy punches were used to create reservoirs and cell seeding pores. Glass coverslips (VWR) were cleaned with 0.2 M HCl overnight, rinsed with water and isopropanol, and dried with compressed air.

##### Covalent bonding protocol

For live cell imaging experiments, the migration devices were assembled after plasma treatment of both the PDMS blocks and glass coverslips for 5 minutes, by gently pressing the PDMS blocks on the activated coverslips to form covalent bonds. The assembled devices were heated on a hot plate at 95°C for 5 minutes to improve adhesion.

##### Non-covalent bonding protocol

For fixed cell IF staining experiments, the migration devices were assembled after plasma treatment of only the glass coverslips for 5 minutes, by gently pressing the PDMS blocks on the activated coverslips to form non-covalent bonds. The assembled devices were heated on a hot plate at 65°C for 1 hour to improve adhesion, and then plasma treated for another 5 minutes after prolonged (10 minutes) vacuuming.

After bonding, devices were filled with 70% ethanol (the step was skipped in *non-covalent bonding protocol*) to sterilize, then rinsed and coated with 50 µg/mL rat tail type-I collagen (Corning) in 0.02 N acetic acid (Sigma) overnight at 4°C. After coating, the devices were rinsed with complete DMEM before cell seeding. For overnight live cell imaging experiments and standard IF staining experiments, 30,000 cells suspended in 5 µl of media were seeded into individual loading chambers of the migration devices. A 0-to-200 ng/mL platelet-derived growth factor (PDGF, Cell Signaling) gradient in DMEM was established for migration of human fibroblasts. For low confluency IF staining experiments (5-days-long time-lapse imaging, staining with inhibitor treatments and siRNA depletions), 6,000 cells were seeded, and a 2-to-10% (v/v) FBS gradient in DMEM was established for increased cell migration capacity. Subsequently, devices were put in the incubator, for either a minimum of 6 hours to allow cell adhesion before live cell imaging under a LSM700 confocal microscopy (Zeiss), or around 48 hours to allow cell migration before fixation and IF staining. For live cell imaging, cells were cultured and imaged in FluoroBrite DMEM (Gibco) supplemented with 10% FBS, 1% Pen-Strep, GlutaMax (Gibco) and 25 mM HEPES (Gibco). Where indicated, media inside the devices was supplemented with DMSO (vehicle) or inhibitors throughout the experiments. For experiments with GFP-HDAC3, SPY555-DNA (Cytoskeleton, 1:3,000) was used to label nuclear regions.

#### Immunofluorescence staining

For staining of cells in microfluidic migration devices, cells were fixed with 4% paraformaldehyde (PFA, Sigma) in PBS for 30 minutes at room temperature. After PBS washes, the devices were carefully removed by inserting a razor blade from the PDMS edges into the interface between the PDMS and the glass coverslips, and then gradually lifting up the PDMS blocks. For staining of cells seeded directly on glass coverslips, cells were fixed with 4% PFA in PBS for 10 minutes at room temperature. The fixation step was followed by permeabilization with 0.3% Triton-X100 (Sigma) in PBS for 15 minutes, blocking with 3% bovine serum albumin (BSA) (Sigma) in PBS supplemented with 0.2% Triton-X100 and 0.05% Tween-20 (Sigma) for 1 hour at room temperature, and incubation overnight at 4°C with primary antibodies (rabbit anti-H3K9me3, Abcam, ab8898, 1:1,000; rabbit anti-H3K27me3, Millipore, 07-449, 1:250; mouse anti-H3K9ac, GeneTex, GT464, 1:500; rabbit anti-pSer424-HDAC3, Cell Signaling, 3815, 1:200; mouse anti-lamin A/C, Santa Cruz, sc-376248, 1:300; mouse anti-lamin A, Santa Cruz, sc-518013, 1:100; mouse anti-emerin, Leica Biosystems, NCL-EMERIN, 1:200, rabbit anti-pSer2 RNA polymerase II, Abcam, ab5095, 1:500) in the same blocking buffer. The samples were then washed with PBS and stained with Alexa Fluor secondary antibodies (Invitrogen) and DAPI (Sigma, 1:1,000). For staining of 5-methylcytosine, cells were treated with 4N HCl for an hour at 37°C for denaturing of DNA, and washed with PBS before blocking and primary antibody incubation (mouse-anti 5-methylcytosine, Eurogentec, BI-MECY-0100, 1:250). After PBS washes, the samples were mounted with Hydromount Nonfluorescing Mounting Media (Electron Microscopy Sciences) onto glass slides (VWR) for confocal microscopy imaging. Alternatively, cells were permeabilized with 0.2% Triton-X100 (Sigma) in PBS for 10 minutes, and an extra 5% normal donkey serum (Millipore) was added to blocking buffer for certain primary antibodies (mouse anti-HDAC3, Cell Signaling, 3949, 1:250; goat anti-HP1α, Abcam, ab77256, 1:300).

For staining of cells in 3D collagen matrices, cells were fixed with 4% paraformaldehyde (PFA, Sigma) in PBS for 30 minutes at room temperature. The fixation step was followed by permeabilization with 0.5% Triton-X100 (Sigma) in PBS for 25 minutes at room temperature, washing by PBS for 15 mins for 3 times, blocking with 3% bovine serum albumin (BSA) (Sigma) in PBS supplemented with 0.2% Triton-X100, 0.05% Tween-20 (Sigma), and 5% normal donkey serum (Millipore) overnight at room temperature, and incubation for 3 days at 4°C with primary antibodies (rabbit anti-H3K27me3, Millipore, 07-449, 1:150; mouse anti-H3K9ac, GeneTex, GT464, 1:350). The matrices were then washed by PBS for 1 hour for 3 times, followed by a second blocking step overnight at 4°C, and incubation for 1 day at room temperature with Alexa Fluor secondary antibodies (Invitrogen) and DAPI (Sigma, 1:1000). The matrices were then washed by PBS for 1 hour for 3 times before confocal microscopy imaging.

#### 5-ethynyl uridine labeling

To label nascent mRNA, cells were pulsed and labeled as previously described (Jao and Salic, 2008), and following manufacturer’s protocols. Cells migrating in microfluidic devices were pulsed with 1 mM 5-ethynyl uridine (EU, Jena Bioscience) in complete DMEM for 4 hours, and then fixed with 4% PFA in PBS for 30 minutes. After two PBS washes, the devices were carefully removed as described in the IF staining section. The cells were then permeabilized with 0.3% Triton-X100 (Sigma) in PBS for 15 minutes, and incubated for 30 minutes with freshly prepared EU labeling buffer containing 100 mM Tris base (Sigma), 100 μM Alexa Fluor 488 conjugated-azide (Invitrogen), 4 mM CuSO_4_ (Sigma), and 100 mM ascorbic acid (Sigma) in water. After the incubation, cells were washed with PBS, and another round of 30 minutes incubation with EU labeling buffer and PBS washes was repeated. Cells were then incubated with primary antibodies and proceeded with the IF staining protocol.

#### Protein immunoblot analysis

Cells were lysed in RIPA buffer or high-salt RIPA buffer supplemented with cOmplete EDTA-Free protease inhibitor (Roche) and PhosSTOP phosphatase inhibitor (Roche) on ice for 3 minutes. Lysates were then vortexed for 5 minutes, and spun down at 4°C. For lysates blotting for proteins that form network scaffolds (lamin A/C and emerin), lysates were sonicated and briefly boiled before spinning down to remove potential DNA contamination. Supernatant were then transferred to new tubes. Protein concentration was quantified by Bio-Rad Protein Assay and a Model 550 Microplate Reader. 25 µg of protein lysate was heat-denatured at 95°C for 5 minutes in Laemmli sample buffer (Bio-Rad), and then separated using a 4 to 12% Bis-Tris polyacrylamide gel (Invitrogen) following a standard SDS-PAGE protocol. Protein was transferred using a semi-dry system (Bio-Rad) to a polyvinylidene fluoride membrane (Millipore, IPVH00010) for 1.5 hour at a current of 16 mA. Membranes were blocked with 3% BSA in Tris-buffered saline containing 0.1% Tween-20 (TBST) for 1 hour at room temperature, and then incubated overnight at 4°C with primary antibodies (rabbit anti-H3K9me3, Abcam, ab8898, 1:3,000; rabbit anti-H3K27me3, Millipore, 07-449, 1:3,000; mouse anti-H3K9ac, GeneTex, GT464, 1:3,000; mouse anti-lamin A/C, sc-376248, Santa Cruz, 1:1,000; mouse anti-emerin, Leica Biosystems, NCL-EMERIN, 1:500; mouse anti-HDAC3, Santa Cruz, sc-136290, 1:500; mouse anti-CK2α, Santa Cruz, sc-365762, 1:250) in the same blocking buffer. After TBST rinses, membranes were incubated with loading control primary antibody (mouse anti-histone H3, Abcam, ab195277, 1:5,000; rabbit anti-histone H3, Cell Signaling, 4499, 1:1,000; mouse anti-GAPDH, Proteintech, 60004-1-Ig, 1:20,000; rabbit anti-pan-actin, Cell Signaling, 8456, 1:1,000) for 1 hour at room temperature. Protein bands were detected using IRDye 680LT and IRDye 800CW secondary antibodies (LI-COR), imaged with an Odyssey CLx imaging system (LI-COR), and analyzed with Image Studio Lite (LI-COR).

#### Extended imaging using an incubator microscope

Long-term imaging was performed using an IncuCyte Live-Cell Analysis System (Sartorius) inside the cell culture incubator. Cells expressing NLS-GFP and migrating inside the microfluidic migration devices were imaged using the IncuCyte filter module for 5 days, every 10 minutes, with a 20× objective. The acquired migration movies were processed and exported using the IncuCyte ZOOM software (Sartorius). At the end of imaging, the cells were immediately fixed for IF staining of heterochromatin and euchromatin marks.

#### Confocal microscopy

Fixed cells on coverslips and live cells migrating in microfluidic devices were imaged with an inverted Zeiss LSM700 confocal microscope. Z-stack images were collected using 20× air (NA = 0.8) or 40× water-immersion (NA = 1.2) objectives. Airy units for all images were set at 1.0. For overnight live cell imaging, the image acquisition was automated through ZEN (Zeiss) software at 10 minutes intervals, at 37°C in a heated stage chamber for 12-16 hours. Where indicated, temperature of the heated chamber was lowered to 33°C.

#### Reflected light microscopy

Images of collagen matrices were acquired by synchronizing between fluorescence and reflectance tracks on an inverted Zeiss LSM880 confocal microscope, using a 40× water-immersion (NA = 1.2) objective lens. Reflectance was acquired with the 561 nm excitation line, collecting with a 550-570 nm bandwidth.

#### Fluorescence recovery after photobleaching (FRAP)

Images were acquired with an inverted Zeiss LSM700 confocal microscope using a 63× oil-immersion (NA = 1.4) objective, at 37°C in a heated stage chamber. Three 1.8 μm by 1.8 μm regions were selected in each cell: one as the “unbleached” control region, one as the low GFP-HP1α intensity “euchromatin” control region, and one as the high GFP-HP1α intensity “inquiry” region (with GFP-HP1α foci or enrichment). The three regions were first imaged for 2 seconds at 1% power of 488 nm laser and 0.2 second intervals, to calculate the pre-bleach values. The euchromatin control and the inquiry regions were then bleached with a laser pulse of 12% power for 20 iterations. Following bleaching, the three regions were imaged for at least 24 seconds at 1% power and 0.2 second intervals. The FRAP image acquisition was automated through ZEN (Zeiss) software. Fluorescence recovery was measured as percentage of recovery (with pre-bleach value as 100%), normalized to unbleached control. Immobile fractions were measured as fluorescence intensity that had not recovered after 24 seconds, after the recovery curve had reached plateau (by around 22 seconds).

#### Assay for Transposase-Accessible Chromatin using sequencing (ATAC-seq)

To achieve consistent bonding of collagen matrices, 35-mm glass-bottom dishes (World Precision Instruments) were treated with 1% polyethylenimine, followed by 0.1% glutaraldehyde treatment and washed with PBS as described (Cross et al., 2010). To generate single cell-containing collagen matrices at specified collagen concentrations (low: 0.3 mg/mL; medium: 1.0 mg/mL; high: 1.7 mg/mL), acidic solution of rat tail type-I collagen (Corning) was supplemented with complete DMEM and NaOH to reach a neutral pH of 7.4, and then mixed with cell suspension to reach a final density of 100,000 cells per mL, and a volume of 800 μL per dish. Cells were allowed to migrate in the matrices for 48 hours in culture media containing a broad MMP inhibitor, GM6001 (Millipore, 20 μM). Where indicated, vehicle control with DMSO was also used. After migration, single cell-containing collagen was carefully collected by pipette tip into microcentrifuge tubes, and an equal volume of 5,000 U (μmol/min) collagenase type 7 (Worthington) dissolved in DPBS (Gibco) was added into each tube for incubation for 15 minutes at 37°C. After digestion, cells were spun down at 4°C, and then resuspended into freezing media (10% DMSO, 20% FBS, and 70% culture media) for freeze-down. A total of four independent replicates were collected for cells in each collagen concentration. Frozen cells were sent to Cornell Univerisity’s Transcriptional Regulation & @Expression Facility (TREx) for library preparation.

For library preparation of ATAC-seq, viable frozen cells were thawed rapidly at 37°C, and then slowly diluted using DMEM supplemented with 10% FBS until the diluted volume reached at least 10-fold of the initial volume. If needed, samples were concentrated by centrifuging at 500 RCF for 5 minutes, followed by removal of supernatant. Omni-ATAC protocol (Corces et al., 2017) was followed, with the following modifications: a cell count of up to 25,000 per sample was used, and cells were washed twice in ATAC-RSB (resuspension buffer) containing 0.1% Tween-20 prior to lysis and transposition. A reaction volume of 25 μL was used for transposition. After the transposition, half of the reaction product was amplified by PCR using a total of 11 cycles without qPCR testing, followed by SPRIselect bead (Beckman Coulter) clean-up (2:1 bead ratio). All the buffers (RSB-Tween, lysis buffer, and TD buffer) were made fresh before use. After the library preparation, samples were quantified by a Qubit (Thermo Fisher) and the size distribution was determined by a Fragment Analyzer (Agilent). If necessary, a digital PCR assay (Bio-Rad QX200) was used to accurately quantify the sequenceable molarity. The libraries were then sequenced on an Illumina HiSeqX at Novogene with 2×150 paired-end reads (Corces et al., 2017).

ATAC-seq data quality was assessed with fastqc (Andrews, 2010). Adapter sequences were trimmed using fastp (Chen et al., 2018). Reads were aligned to the hg38 reference genome assembly using bowtie2 (Langmead and Salzberg, 2012) in sensitive-local mode. Alignments were filtered to remove reads with multiple valid alignments (mapq > 1). Coverage signal tracks were generated using deepTools bamCoverage (Ramírez et al., 2014) and normalized on a per-million basis using R(R Core Team, 2020). Peaks were called using macs2 (Zhang et al., 2008) with default settings, pooling all datasets as input, using a minimum -log10 q-value threshold of 100. Reads mapping within peaks were counted using featureCounts (Liao et al., 2014), and differential accessibility was determined using DESeq2 (Love et al., 2014), considering peaks with adjusted p-values lower than 0.1 as significantly differentially accessible (DA). All DESeq2 comparisons were performed using the full matrix of high, medium, and low concentration collagen ATAC-seq scores, so pairwise comparisons were assessed for significance in the context of global variation across all three conditions. HOMER (Hypergeometric Optimization of Motif EnRichment) (Heinz et al., 2010) was used for annotation of DA peaks. Metaprofiles were generated using the BRGenomics R package (DeBerardine, 2020), and are representative of average and 75% confidence interval over indicated sets of peaks calculated by subsampling 10% of peaks 1,000 times. Peaks that overlap with the region from -1,000 to transcription start site (TSS) of any annotated transcript in the hg38 genome assembly were considered promoter-associated, and all other peaks were considered non-promoter-associated. All promoter-associated peaks with the nearest non-promoter-associated peaks closer than 500 kb were considered enhancer-promoter interactions (EPI). Enrichment of transposable element (TE) overlaps with the DA peaks was performed using the script TE-anaysis_Shuffle_bed.pl (https://github.com/4ureliek/TEanalysis) (Kapusta et al., 2013; Lynch et al., 2015). Scripts used to perform these analyses including exact parameters can be accessed at https://github.com/jaj256/ConfinedATAC.

Genomic Regions Enrichment of Annotations Tool (GREAT) (McLean et al., 2010) was used to plot each DA peak’s distance to the TSS of its associated genes, and to analyze enrichment of Gene Ontology (GO) Biological Process and Cellular Component terms. Ingenuity Pathway Analysis (IPA, Qiagen, https://www.qiagenbioinformatics.com/products/ingenuitypathway-analysis) (Krämer et al., 2014) was used to analyze enrichment of canonical pathways, and to generate predicted regulatory networks. DREME (Bailey, 2011) was used for motif analysis, and TomTom (Gupta et al., 2007) was used to match known motifs of transcription factors to the enriched motifs. UCSC genome browser (http://genome.ucsc.edu) (Kent et al., 2002), ENCODE annotation data (Rosenbloom et al., 2013), and GENCODE annotation data (Harrow et al., 2012) were used to generate screenshots of EPIs.

The distance of DA peaks to centromeres or telomeres were calculated by using the ‘findOverlapsOfPeaks’ function from the ChIPpeakAnno package (Zhu et al., 2010). The positions of the centromeres were obtained from the Bioconductor website (https://rdrr.io/bioc/rCGH/man/hg38.html). The positions of the telomeres were obtained from the Bioinformatics Work Notes website (https://blog.gene-test.com/telomeric-regions-of-the-human-genome/).

For immunofluorescence staining validation, single cells were embedded into collagen matrices and allowed 48 hr of migration in the matrices, as described in the previous paragraphs. After collagenase digestion and collection, cells were seeded onto coverslips coated with 0.01% of poly-L-Lysine (Sigma). After 45 minutes, cells were fixed and stained as described in the “immunofluorescence staining” section, except that the blocking buffer used 5% donkey serum, and a second blocking step of 20 minutes was added before incubation with secondary antibodies.

#### RNA extraction and quantitative PCR (qPCR)

To achieve consistent bonding of collagen matrices, 35-mm glass-bottom dishes (World Precision Instruments) were treated with PEI and glutaraldehyde, as described in the ATAC-seq section. To generate single cell-containing collagen matrices at specified collagen concentrations (low: 0.3 mg/mL; high: 1.7 mg/mL), acidic solution of rat tail type-I collagen (Corning) was supplemented with complete DMEM and NaOH to reach a neutral pH of 7.4, and then mixed with cell suspension to reach a final density of 100,000 cells per mL, and a volume of 1.6 mL per dish. Cells were allowed to migrate in the matrices for 48 hours in culture media containing a broad MMP inhibitor, GM6001 (Millipore, 20 μM). After migration, each dish was rinsed with diethylpyrocarbonate (DPEC, Sigma)-treated DPBS (Gibco), and single cell-containing collagen was carefully collected by pipette tip into microcentrifuge tubes, and an equal volume of 5,000 U (μmol/min) collagenase type 7 (Worthington) dissolved in DPBS (Gibco) was added into each tube for incubation for 15 minutes at 37°C. After digestion, cells were spun down at 4°C, and total RNA was extracted using RNeasy Plus Mini Kit (Qiagen). Genomic DNA was removed using the TURBO DNA-free Kit (Thermo Fisher), and mRNA was converted to cDNA with the iScript cDNA Synthesis Kit (Bio-Rad) for a final volume of 20 uL per sample, following the manufacturers’ instructions. A total of six replicates from two independent experiments were collected for cells in high or low concentration collagen for qPCR analysis. qPCR was performed on a LightCycler 480 qPCR system (Roche), using the LightCycler 480 SYBR Green I Master mix (Roche). Gene expression level of *HDAC3* and *CBX5/HP1α* were probed using the primer sequences listed in Table S6; Geometric mean of *β-actin, GAPDH* and *18S* gene expression levels were used as housekeeping controls. The Ct (cycle threshold) values were calculated using the second derivative max method on the LightCycler 480 Software (Roche). The primers (Integrated DNA Technologies) used for each of these genes are listed in Table S6. qPCR cycles were performed using the settings listed in Table S7.

#### Image analysis

Image sequences were analyzed using ZEN (Zeiss), ImageJ or MATLAB (Mathworks) using only linear intensity adjustments uniformly applied to the entire image region. Region-of-interest intensities were extracted using ZEN or ImageJ. All confocal image stacks were three-dimensionally reconstructed as maximum intensity projections, and then processed with standard background subtraction (rolling ball). To quantify normalized heterochromatin values, nuclear areas were selected by thresholding using the median-filtered DAPI channel. Mean grey intensity of the heterochromatin mark (H3K9me3 or H3K27me3) and the euchromatin mark (H3K9ac) were quantified, and the heterochromatin intensity was divided by the euchromatin intensity to obtain normalized heterochromatin values. Apoptotic fragments and overlapped nuclei were manually excluded from the dataset. Experiments with high variability of cell seeding between control and experimental groups were excluded. For all other quantification of staining intensities, nuclear areas were selected by thresholding using the median-filtered DAPI channel (or comparable nuclear signal when DAPI was not available), and mean grey intensities of channels of interests were quantified. All staining thresholding and quantification were performed through custom ImageJ Macros (available at https://github.com/chiehrenhsia/ConfinedMigration) to ensure consistency and reproducibility. For generation of normalized heterochromatin images, the heterochromatin mark channel (H3K9me3 or H3K27me3) was divided by the euchromatin mark (H3K9ac) channel using the “image calculator” function in ImageJ. In the euchromatin (H3K9ac) channel, pixels with 0 value were converted to 1 to ensure division without errors. The normalized image was then median-filtered and a nuclear mask of median-filtered DAPI channel was applied to exclude artifacts of non-nuclear background noise. To quantify cells with persistent GFP-HP1α enrichments, time-lapse cell migration movies were blinded to the observer to record the time duration of GFP-HP1α enrichment formation after the start of nuclear transit in Rainbow RGB pseudo-color channel. An enrichment was defined by at least 30% increase in nuclear bleb GFP intensity compared to the main nucleus. The durations of all GFP-HP1α enrichments were plotted, and enrichments with durations longer than the mean + 95% confidence interval were considered significantly longer than the mean, therefore defined as persistent enrichments. To quantify GFP-HDAC3 nucleoplasmic-to-cytoplasmic (Nuc/Cyto) ratio, nuclear region was selected using SPY555-DNA signal, and whole-cell region was traced manually in ImageJ. Cytoplasmic GFP signal was calculated by whole-cell GFP signal subtracting nuclear GFP signal. Nuc/Cyto ratio was then calculated by dividing nuclear signal with cytoplasmic signal. Nuc/Cyto ratio of GFP-HDAC3 from 2-6 time points immediately before and after nuclear transit were quantified and averaged. To quantify nuclear transit time through microfluidic migration channels, time-lapse cell migration movies were analyzed using a custom MATLAB program, as described previously(Elacqua et al., 2018). Any single outlier with a transit time 180 minutes greater than the rest of the dataset was excluded in each experiment. Graphs were generated in Prism 8.2.0 or 9.3.1 (GraphPad) or Excel (Microsoft), and figures were assembled in Illustrator (Adobe).

### Quantification and Statistical Analysis

Unless otherwise noted, all experimental results are pooled from at least three independent experiments. For data with normal distribution, two-tailed Student’s *t* test with Welch’s correction (comparing one variable in two conditions), or one-way analysis of variance (ANOVA) with Tukey’s post hoc test for multiple comparisons (comparing one variable in more than two conditions) was used. For data where two factors were involved, two-way ANOVA with Tukey’s post hoc test for multiple comparisons was used. For data with non-normal distribution, Wilcoxon Rank Sum test (comparing one variable in two conditions), or Kruskal-Wallis H test with Dunn’s post hoc test for multiple comparisons (when two factors were involved) was used. For paired data (Nuc/Cyto ratio of GFP-HDAC3 before and after nuclear transit), paired *t* test was used. All statistical tests were performed using Prism version 9.3.1 for Mac, or version 8.2.0 for Windows (GraphPad). Statistical details are provided in the figure legends. Unless otherwise noted, error bars represent the standard error of the mean (SEM).

## Acknowledgments

The authors thank Cornell University’s Transcriptional Regulation & Expression Facility (TREx), especially Jen Grenier, for ATAC-seq library preparation their help with the analysis; Cornell University’s Biotechnology Resource Center (BRC) Genomics Facility for ATAC-seq library quality control and sequencing; Cornell University’s Statistical Consulting Unit (CSCU) for suggestions and reviewing of statistical analysis. The authors thank Philipp Isermann, Tyler Kirby, Pragya Shah, and other members of the Lammerding group for their helpful discussions and support. This work was supported by awards from the National Institutes of Health (R01 HL082792, R01 GM137605, U54 CA210184 to J.L.), the Department of Defense Breast Cancer Research Program (Breakthrough Award BC150580 to J.L.), the National Science Foundation (CAREER Award CBET-1254846 to J.L.), the Volkswagen Stiftung (to J.L.) and the Taiwanese Ministry of Education (GSSA-2017 to C.-R.H.). J.J. was supported by National Human Genome Research Institute fellowship (F31HG010820). The content is solely the responsibility of the authors and does not necessarily represent the official views of the National Institutes of Health. This work was performed in part at the Cornell NanoScale Science & Technology Facility (CNF), a member of the National Nanotechnology Coordinated Infrastructure NNCI), which is supported by the National Science Foundation (Grant NNCI-2025233). This work was performed in part using the Zeiss LSM880 microscope at the Cornell Biotechnology Resource Center, which is supported by the National Institutes of Health (S10 OD018516).

## Author Contributions

C.-R.H. and J.L. conceptualized and designed all experiments; C.-R.H., J.M., and O.H. performed experiments; J.J. and S.L. optimized the ATAC-seq analysis pipeline and performed the ATAC-seq analysis; R.A. designed and manufactured the 10-μm tall microfluidic migration devices; C.-Y.C. optimized initial histone modifying enzyme inhibitor conditions; C.-R.H. and J.L. wrote the paper; all authors contributed to the editing of the manuscript; P.S. and J.L. acquired funding.

