## Supplementary Text + Figures for "Confined Migration Induces Heterochromatin Formation and Alters Chromatin Accessibility"

**for**

### Supplemental Figures

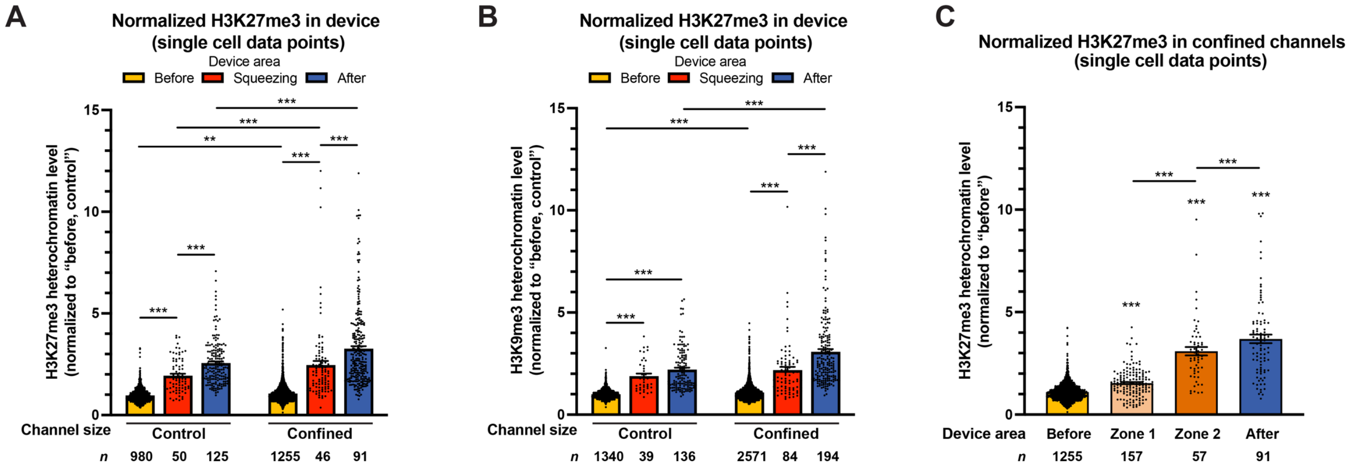


**Figure S1. Confined migration induces heterochromatin formation in cells migrating through microfluidic devices (single cell data points). (A)** . Quantification of normalized H3K27me3 heterochromatin level in HT1080 cells migrating in migration devices. All values are normalized to control channels "before" cells. Data corresponds to results in Figure 1D, but shows individual cell data points. $^{**}p < 0.01$, $^{***}p < 0.001$, two-way ANOVA with Tukey's multiple comparison test. **(B)** Quantification of normalized H3K9me3 heterochromatin level in HT1080 cells migrating in migration devices. All values are normalized to control channels "before" cells. Data corresponds to results in Figure 1F, but shows individual cell data points. $^{***}p < 0.001$, two-way ANOVA with Tukey's multiple comparison test. **(C)** Comparison between normalized H3K27me3 heterochromatin level of HT1080 cells in each area of the device. All values are normalized to "before" cells. Data corresponds to results in Figure 1G, but shows individual cell data points. $^{***}p < 0.001$, one-way ANOVA with Tukey's multiple comparison test. Data are presented as mean ± SEM, from *n* cells (listed in each graph) pooled from three biological replicates.

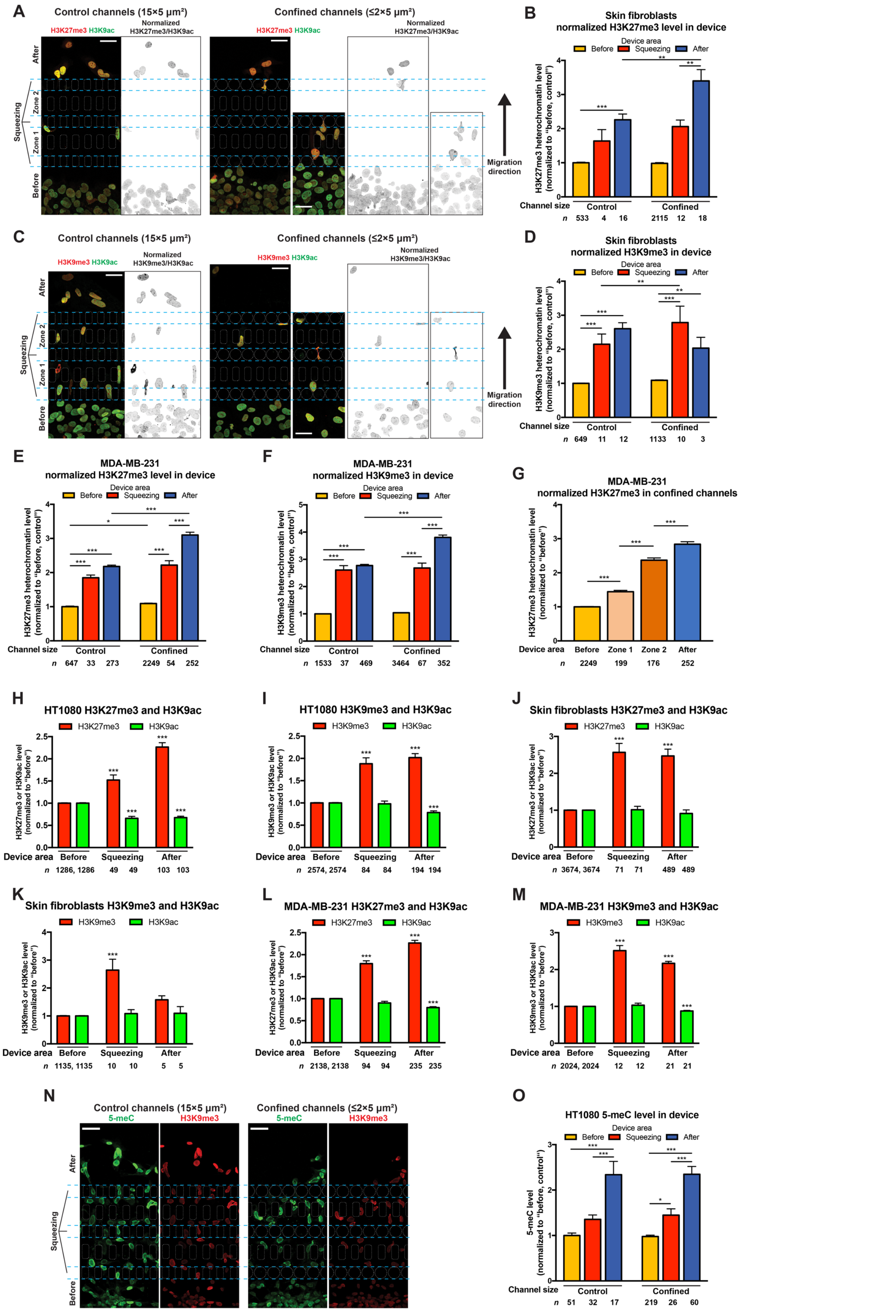


**Figure S2: Confined migration induces heterochromatin formation in various cell types in microfluidic migration devices. (A)** Representative staining of H3K27me3 (red) and H3K9ac (green) in skin fibroblasts migrating in a migration device, in control or confined channels. Normalized heterochromatin level (H3K27me3 divided by H3K9ac) is shown in inverted grayscale. Scale bars: 40 µm. **(B)** Quantification of normalized H3K27me3 heterochromatin level in skin fibroblasts migrating in migration devices. All values are normalized to control channels "before" cells. $^{**}p < 0.01$, $^{***}p < 0.001$, two-way ANOVA with Tukey's multiple comparison test. **(C)** Representative staining of H3K9me3 (red) and H3K9ac (green) in skin fibroblasts migrating in a migration device, in control or confined channels. Normalized heterochromatin level (H3K9me3 divided by H3K9ac) is shown in inverted grayscale. Scale bars: 40 µm. **(D)** Quantification of normalized H3K9me3 heterochromatin level in skin fibroblasts migrating in migration devices. All values are normalized to control channels "before" cells. $^{**}p < 0.01$, $^{***}p < 0.001$, two-way ANOVA with Tukey's multiple comparison test. **(E-F)** Quantification of normalized H3K27me3 and H3K9me3 heterochromatin level in MDA-MB-231 cells migrating in migration devices. All values are normalized to control channels "before" cells. $^{*}p < 0.05$, $^{***}p < 0.001$, two-way ANOVA with Tukey's multiple comparison test. **(G)** Comparison between normalized H3K27me3 heterochromatin level of MDA-MB-231 cells in each area of the migration device. All values are normalized to "before" cells. $^{***}p < 0.001$, one-way ANOVA with Tukey's multiple comparison test. **(H-I)** Quantification of H3K27me3 or H3K9me3 (red), and H3K9ac (green) intensities of HT1080 cells migrating through confined channels in migration devices. All mark levels are normalized to their levels in "before" cells, respectively. $^{***}p < 0.001$, two-way ANOVA with Tukey's multiple comparison test. **(J-K)** Quantification of H3K27me3 or H3K9me3 (red), and H3K9ac (green) intensities of skin fibroblasts migrating through confined channels in migration devices. All mark levels are normalized to their levels in "before" cells, respectively. $^{***}p < 0.001$, two-way ANOVA with Tukey's multiple comparison test. **(L-M)** Quantification of H3K27me3 or H3K9me3 (red), and H3K9ac (green) intensities of MDA-MB-231 cells migrating through confined channels in migration devices. All mark levels are normalized to their levels in "before" cells, respectively. $^{***}p < 0.001$, two-way ANOVA with Tukey's multiple comparison test. **(N)** Representative staining of 5-methylcytosine (5-meC, green) and H3K9me3 (red) HT1080 cells migrating in a migration device, in control or confined channels. Scale bars: 40 µm. **(O)** Quantification of 5-meC intensities in HT1080 cells migrating in migration devices. All values are normalized to control channels "before" cells. $^{*}p < 0.05$, $^{***}p < 0.001$, two-way ANOVA with Tukey's multiple comparison test. Data are presented as mean ± SEM, based on *n* cells (listed in each graph) pooled from three biological replicates, except for data in (O), which are based on two biological replicates.

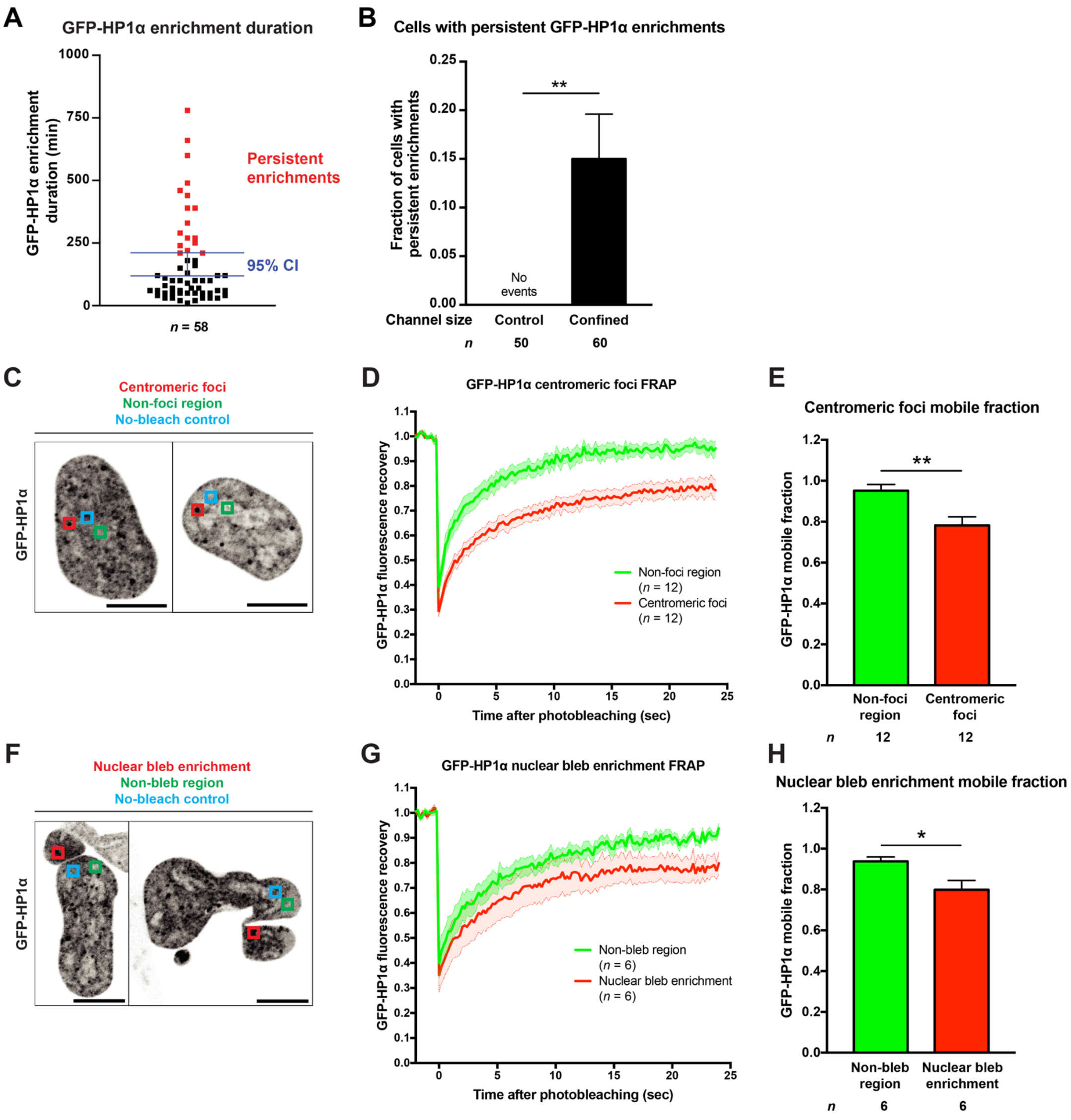


**Figure S3: Nuclear transit induces persistent GFP-HP1α enrichments. (A)** Quantification of GFP-HP1α local enrichment durations in HT1080 cells. Persistent enrichments (red) are defined as having duration at above mean +95% confidence interval (CI). **(B)** Quantification of cells with persistent local enrichment of GFP-HP1α in control channels and confined channels. $^{**}p < 0.01$, student's $t$ test with Welch's correction for unequal variances. **(C)** Representative inverted grayscale image of HT1080 cells with GFP-HP1α centromeric foci (red square) for FRAP analysis. FRAP signal was compared to a non-foci region (green square) and normalized to a no-bleach area (cyan square). Scale bars: 10 µm. **(D)** FRAP curves of HT1080 cells with GFP-HP1α centromeric foci and non-foci regions. Curves and shaded areas represent mean ± SEM. **(E)** Quantification of the mobile fractions (recovered fraction) of GFP-HP1α in non-foci regions and centromeric foci at 24 seconds of recovery. $^{**}p < 0.01$, student's $t$ test with Welch's correction for unequal variances. **(F)** Representative inverted grayscale image of HT1080 cells with GFP-HP1α nuclear bleb enrichments (red square) for FRAP analysis. FRAP signal was compared to a non-bleb region (green square) and normalized to a no-bleach area (cyan square). Scale bars: 10 µm. **(G)** FRAP curves of HT1080 cells with GFP-HP1α nuclear bleb enrichment and non-bleb regions. Curves and shaded areas represent mean ± SEM. **(H)** Quantification of the mobile fractions (recovered fraction) of GFP-HP1α in non-bleb regions and nuclear bleb enrichments at 24 seconds of recovery. $^{*}p < 0.05$, student's $t$ test with Welch's correction for unequal variances. Data are presented as mean ± SEM, based on $n$ cells (listed in each graph) pooled from three biological replicates.

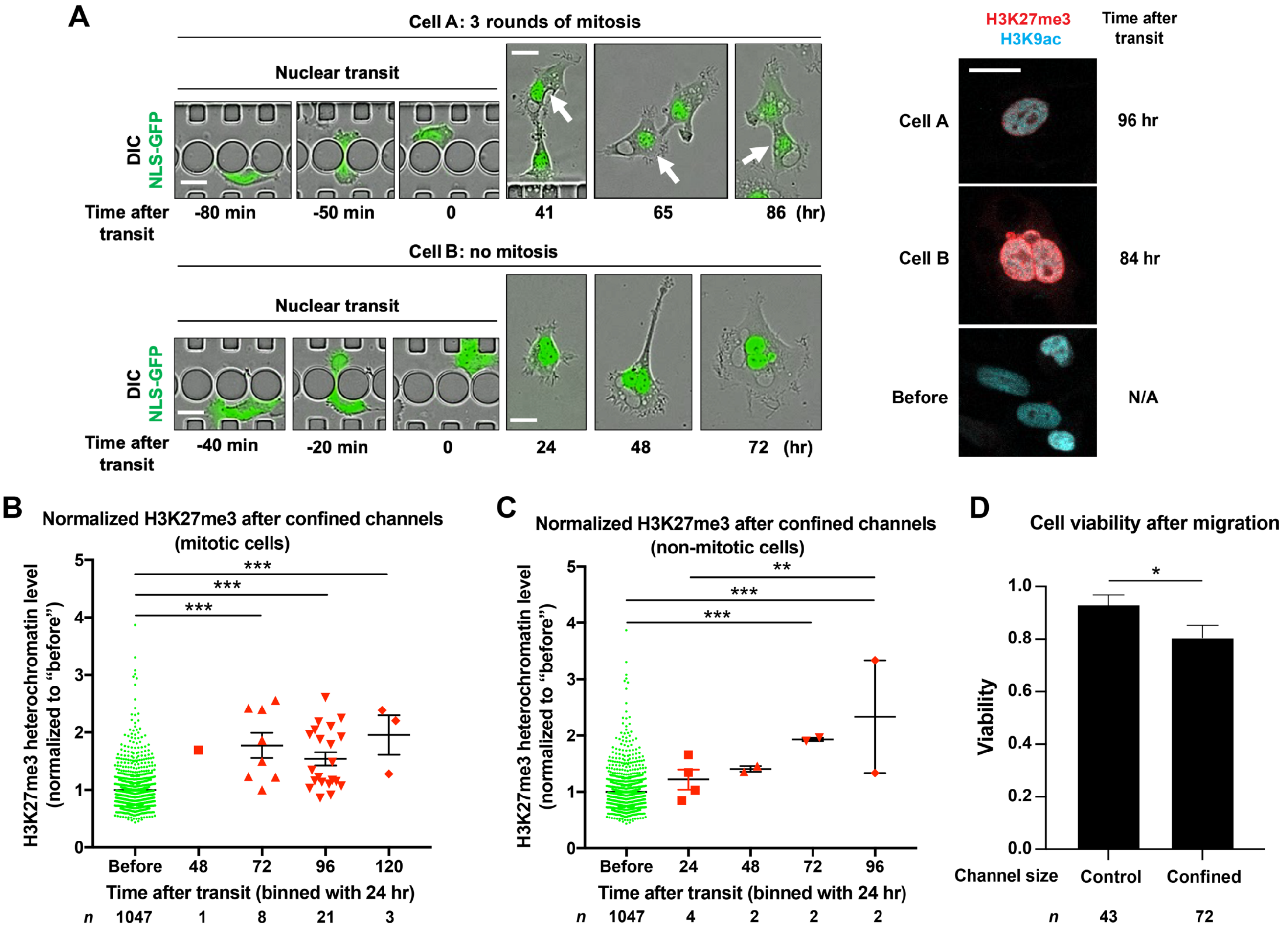


**Figure S4: Confined migration induces persistent heterochromatin formation lasting for at least 5 days. (A)** Representative image sequences of extended live imaging of HT1080 cell migration (left panel), and the chromatin marks staining at fixation (right panel). Green: NLS-GFP; Red: H3K27me3; Cyan: H3K9ac. Scale bars: 20 µm. Cell A is an example of a cell undergoing multiple rounds of mitosis after nuclear transit, while cell B is an example of cells remaining in interphase. Arrows indicate daughter cells; N/A: not applicable (cells did not transit through the channels). Please see also Video S2. **(B)** The correlation of normalized H3K27me3 heterochromatin level with time after the last nuclear transit through confined constrictions in HT1080 cells with mitosis, binned by an interval of 24 hours. All values are normalized to "before" cells. ***$p < 0.001$, one-way ANOVA with Tukey's multiple comparison test. **(C)** The correlation of normalized H3K27me3 heterochromatin level with time after the last nuclear transit through confined constrictions in HT1080 cells without any mitosis, binned by an interval of 24 hours. All values are normalized to "before" cells. **$p < 0.01$, ***$p < 0.001$, one-way ANOVA with Tukey's multiple comparison test. **(D)** Quantification of cell viability of cells that successfully migrated through either the control or confined channels and were located in the "after" region of the device, based on timelapse videos collected over the entire 5-day period of the experiment. *$p < 0.05$, student's $t$ test with Welch's correction for unequal variances. Data are presented as mean ± SEM, based on $n$ cells (listed in each graph) pooled from three biological replicates.

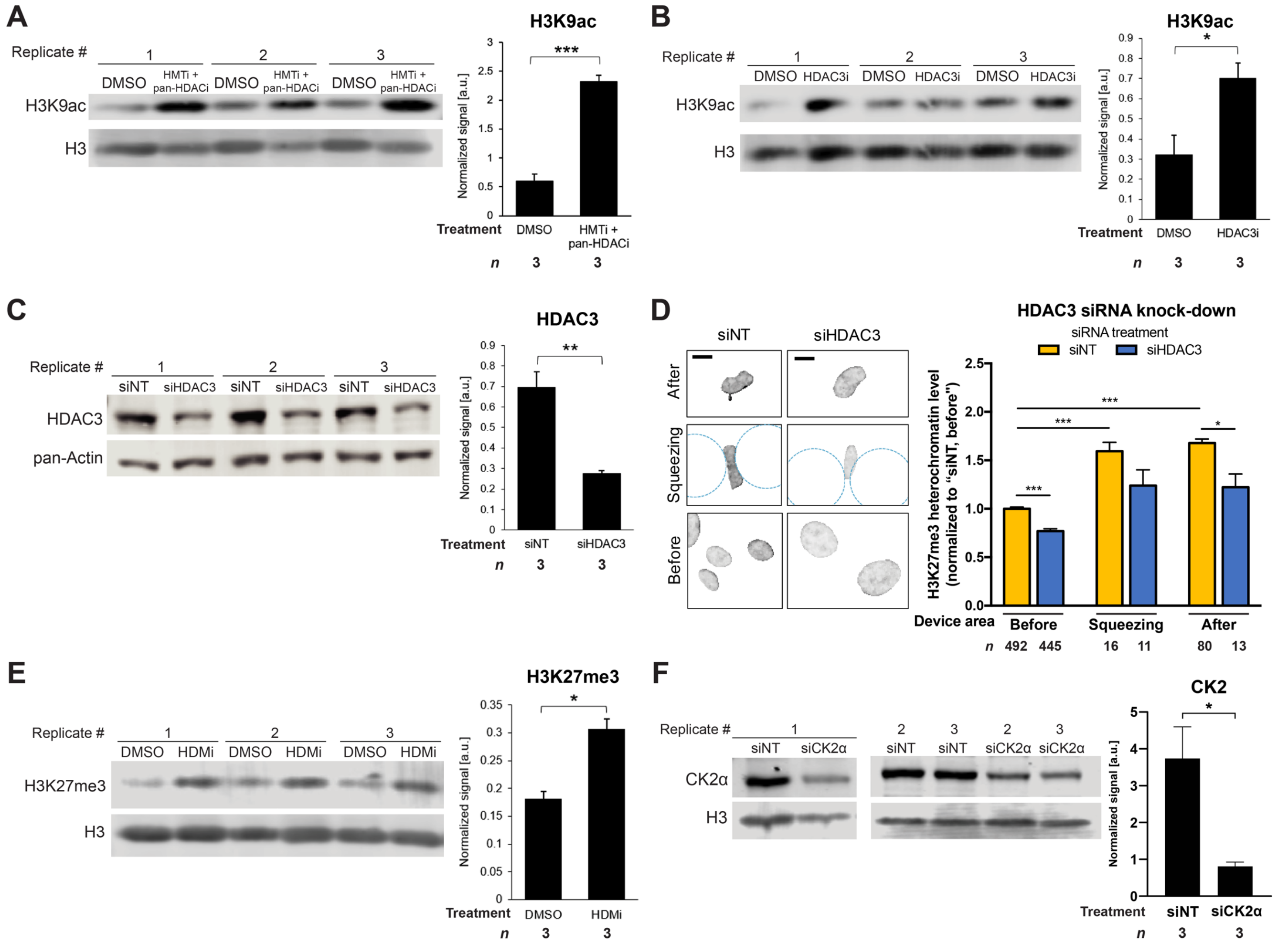


**Figure S5: Confined migration-induced heterochromatin formation is dependent on HDAC3, and the validation of histone modifying enzyme inhibitors. (A)** Validation of HMTi and pan-HDACi treatment on H3K9ac levels compared to DMSO (vehicle) control in HT1080 cells using Western blot. H3K9ac levels are normalized to H3 loading control. ***$p < 0.001$, student's *t* test with Welch's correction for unequal variances. **(B)** Validation of HDAC3i treatment on H3K9ac levels compared to DMSO (vehicle) control in HT1080 cells using Western blot. H3K9ac levels are normalized to H3 loading control. *$p < 0.05$, student's *t* test with Welch's correction for unequal variances. **(C)** Validation of siRNA depletion of HDAC3 (siHDAC3) on HDAC3 levels compared to non-target siRNA (siNT) control in HT1080 cells using Western blot. HDAC3 levels are normalized to pan-actin loading control. **$p < 0.01$, student's *t* test with Welch's correction for unequal variances. **(D)** Left panel: Representative images of normalized H3K27me3 heterochromatin in HT1080 cells treated with non-target siRNA (siNT) or HDAC3 siRNA (siHDAC3) during confined migration. Scale bars: 10 µm. Right panel: Quantification of normalized H3K27me3 heterochromatin in cells treated with siNT or siHDAC3 during confined migration. All values are normalized to siNT "before" cells. *$p < 0.05$, ***$p < 0.001$, two-way ANOVA with Tukey's multiple comparison test. **(E)** Validation of HDMi treatment on H3K27me3 levels compared to DMSO (vehicle) control in HT1080 cells using Western blot. H3K27me3 levels are normalized to H3 loading control. *$p < 0.05$, student's *t* test with Welch's correction for unequal variances. **(F)** Validation of siRNA depletion of CK2 (siCK2α) on CK2 levels compared to non-target siRNA (siNT) control in HT1080 cells using Western blot. CK2 levels are normalized to H3 loading control. *$p < 0.05$, student's *t* test with Welch's correction for unequal variances. Data are presented as mean ± SEM, based on *n* cells or *n* protein lysates (for Western blots) pooled from at least three biological replicates.

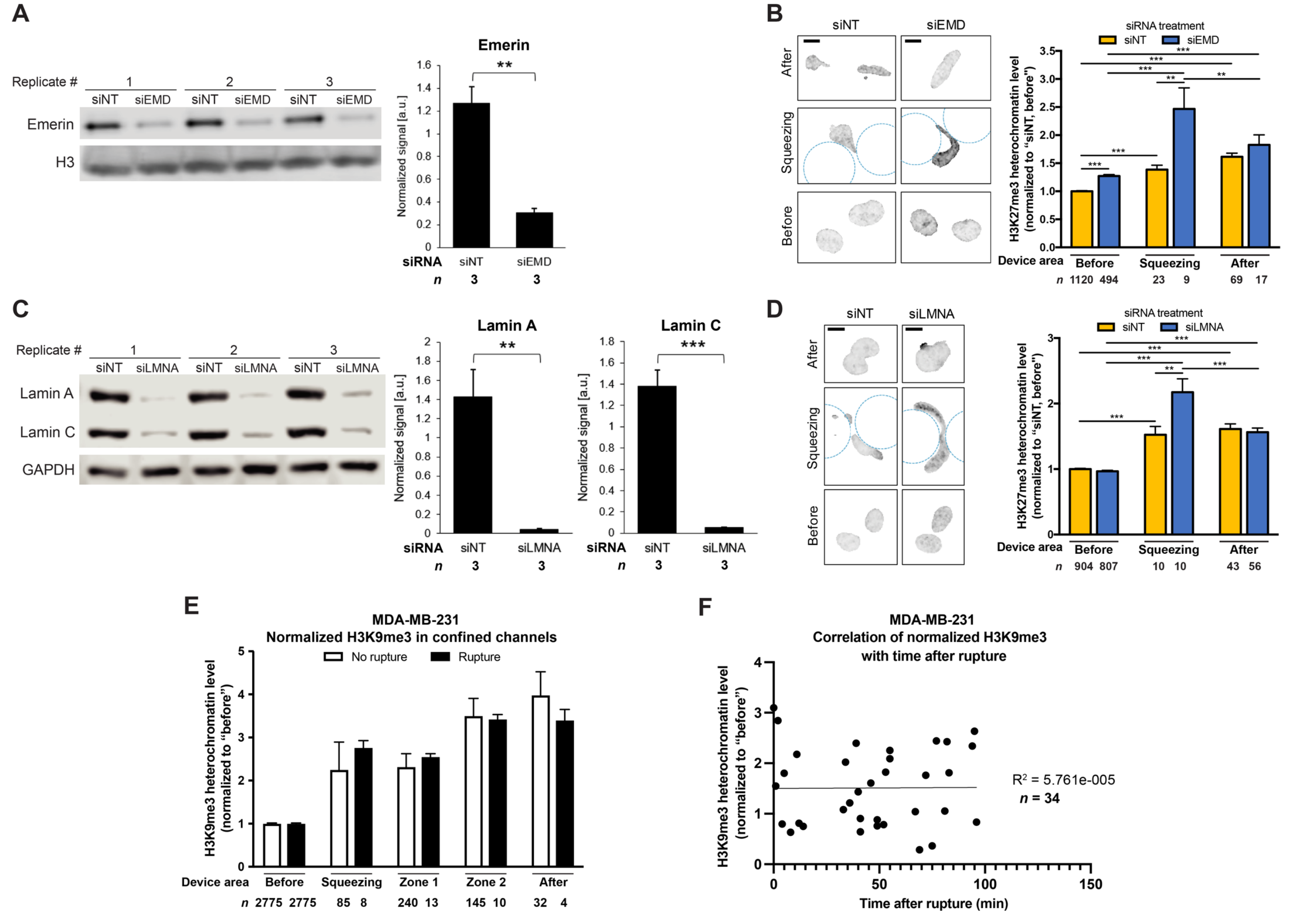


**Figure S6: Confined migration-induced heterochromatin formation is affected by lamin A/C and emerin, but independent of nuclear envelope rupture.** **(A)** Validation of siRNA depletion of emerin (siEMD) on emerin levels compared to non-target siRNA (siNT) control in HT1080 cells using Western blot. Emerin levels are normalized to H3 loading control. $^{**}p < 0.01$, student's $t$ test with Welch's correction for unequal variances. **(B)** Left panel: Representative images of normalized H3K27me3 heterochromatin in HT1080 cells treated with non-target siRNA (siNT) or emerin siRNA (siEMD) during confined migration. Scale bars: 10 µm. Right panel: Quantification of normalized H3K27me3 heterochromatin in cells treated with siNT or siEMD during confined migration. All values are normalized to siNT "before" cells. $^{**}p < 0.01$, $^{***}p < 0.001$, two-way ANOVA with Tukey's multiple comparison test. **(C)** Validation of siRNA depletion of lamin A and C (siLMNA) on lamin A and C levels compared to non-target siRNA (siNT) control in HT1080 cells using Western blot. Lamin A and C levels are normalized to GAPDH loading control. $^{**}p < 0.01$, $^{***}p < 0.001$, student's $t$ test with Welch's correction for unequal variances. **(D)** Left panel: Representative images of normalized H3K27me3 heterochromatin in HT1080 cells treated with non-target siRNA (siNT) or lamin A/C siRNA (siLMNA) during confined migration. Scale bars: 10 µm. Right panel: Quantification of normalized H3K27me3 heterochromatin in cells treated with siNT or siLMNA during confined migration. All values are normalized to siNT "before" cells. $^{**}p < 0.01$, $^{***}p < 0.001$, two-way ANOVA with Tukey's multiple comparison test. **(E)** Quantification of normalized H3K9me3 heterochromatin levels in MDA-MB-231 cells experienced NE rupture (orange) or not (blue) during confined migration. All "rupture" and "no rupture" heterochromatin levels are normalized the same level in "before" cells. **(F)** Correlation and linear regression of normalized H3K9me3 heterochromatin levels with time after rupture in MDA-MB-231 cells. All values are normalized to "before" cells. Data are presented as mean ± SEM, based on $n$ cells or $n$ protein lysates (for Western blots) pooled from at least three biological replicates, except for data in (E-F), which are based on two biological replicates.

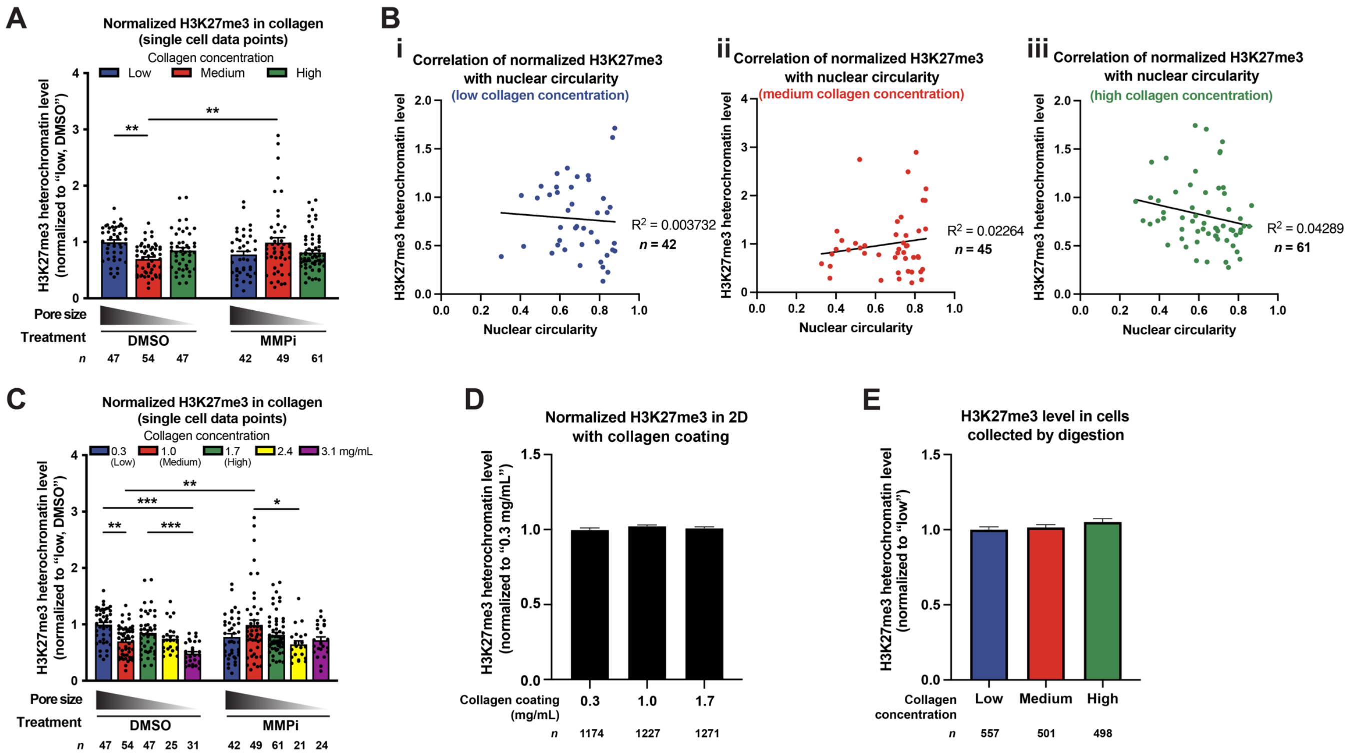


**Figure S7: Confined migration in 3D collagen matrices does not induce global heterochromatin level changes. (A)** Quantification of normalized H3K27me3 heterochromatin level in HT1080 cells migrating in collagen matrices, under DMSO or MMPi treatment. All values are normalized to DMSO-treated cells in low concentration collagen. Data corresponds to results in Figure 5B, but shows individual cell data points. **$p < 0.01$, two-way ANOVA with Tukey's multiple comparison test. **(B)** Correlation and linear regression of normalized H3K27me3 heterochromatin levels with nuclear circularity in HT1080 cells migrating in (i) low, (ii) medium, and (iii) high concentration collagen matrices. All values are normalized to DMSO-treated cells in low concentration collagen. **(C)** Quantification of normalized H3K27me3 heterochromatin level in HT1080 cells migrating in collagen matrices (with two higher concentrations, 2.4 and 3.1 mg/mL), under DMSO or MMPi treatment. All values are normalized to DMSO-treated cells in low concentration collagen. *$p < 0.05$, **$p < 0.01$, ***$p < 0.001$, two-way ANOVA with Tukey's multiple comparison test. **(D)** Quantification of normalized H3K27me3 heterochromatin level in HT1080 cells seeded on 2D coverslips coated with different collagen concentrations. All values are normalized to cells seeded on coverslips coated with 0.3 mg/mL collagen. **(E)** Quantification of normalized H3K27me3 heterochromatin level in HT1080 cells collected from collagen matrices after collagenase digestion. All values are normalized to cells in low concentration collagen. Data are presented as mean ± SEM, based on *n* cells (listed in each graph) pooled from three biological replicates, except for data in (D) and (E), which are based on two biological replicates.

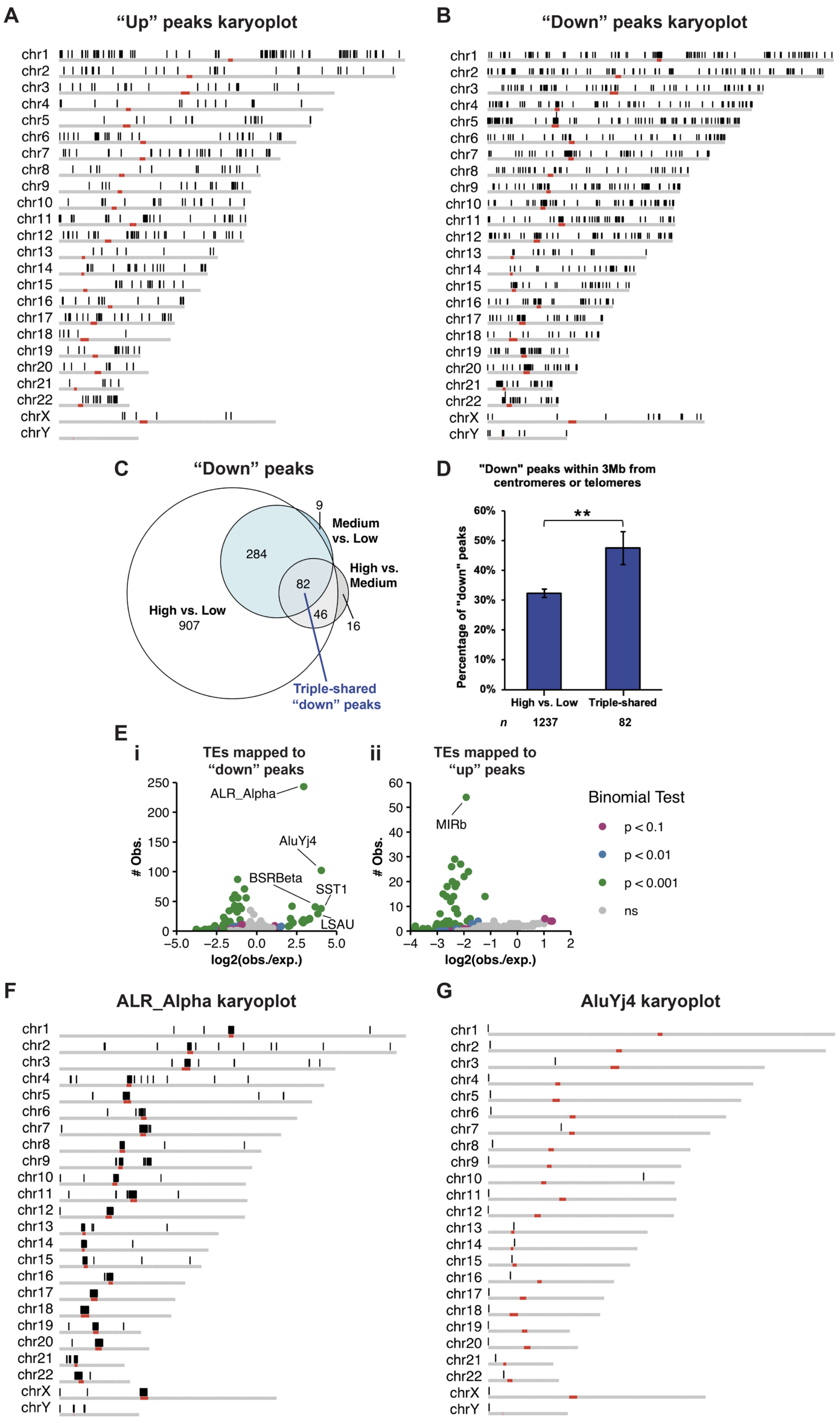


**Figure S8: Confined migration decreases chromatin accessibility at regions near centromeres/telomeres and enriched with transposable elements. (A-B)** Karyoplots of chromosomal locations of "up" and "down" peaks. Red areas: centromeres. Telomeres are located toward the ends of chromosomes. **(C)** Euler diagrams of "down" peaks in all three comparison groups: high versus low, medium versus low, and high versus medium concentration collagen samples. Numbers in each region represent the number of "down" peaks specific to the region. 82 peaks are considered triple-shared "down" peaks among the three comparison groups. **(D)** Quantification of the fraction of peaks that are within 3 Mb from centromeres or telomeres in triple-shared "down" peaks and the rest of the "down" peaks in high versus low concentration collagen samples. $^{**}p < 0.01$, student's $t$ test with Welch's correction for unequal variances. Data are presented as mean ± SEM, based on $n$ DA peaks. **(E)** Binominal tests for enrichment of transposable elements (TEs) mapped to (i) "down" peaks and (ii) "up" peaks, with $p < 0.001$ shown in green. Y-axis: the observed number (# obs.) of TEs mapped to "up" or "down" peaks. X-axis: the log2 ratio of observed versus expected number of TEs. ALR_Alpha and AluYj4 are among the enriched TEs in "down" peaks, while MIRb is among the underrepresented/depleted TEs in "up" peaks. **(F-G)** Karyoplots of chromosomal locations of the enriched TEs, ALR_Alpha and AluYj4. Red areas: centromeres. Telomeres are located toward the ends of chromosomes.

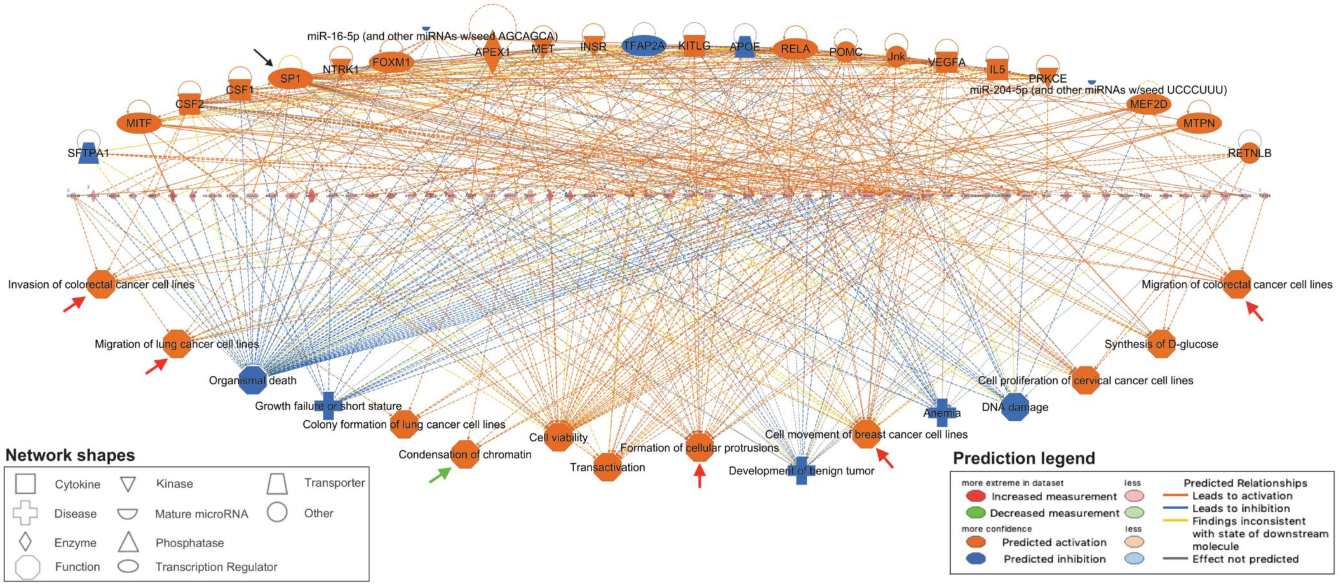


**Figure S9: The full predicted regulation network of "up" peaks-associated genes**. Predicted network of upstream regulators (top) and downstream functions (bottom) of genes associated with "up" peaks (middle), as calculated by IPA (Ingenuity Pathway Analysis). Orange indicates activation, whereas blue indicates inhibition. Black arrow: SP1. Red arrows: activation of cell migration or tumor invasion-related functions. Green arrow: activation of heterochromatin formation-related function.

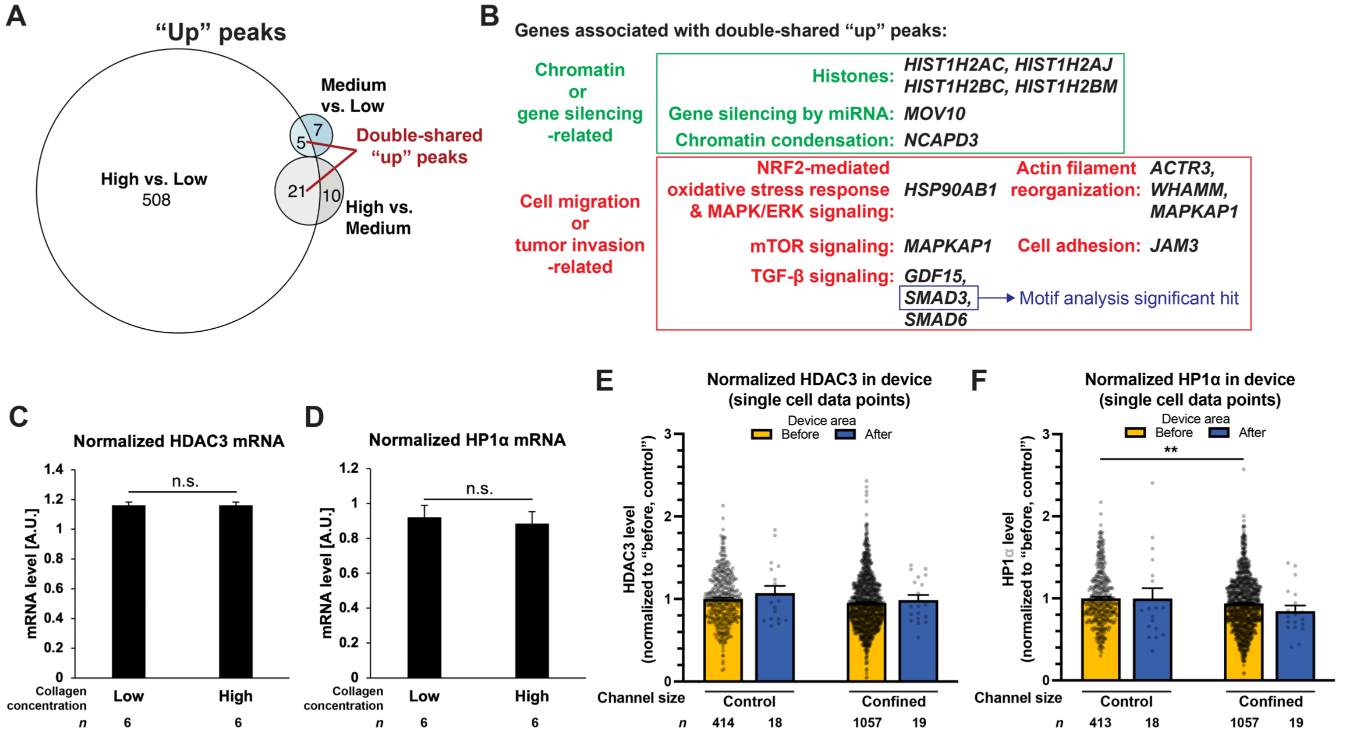


**G** Motif enrichment analysis of "up" peaks

| Motif | Logo | RC Logo | E-value | Unerased E-value | Significant TF hits |
|---|---|---|---|---|---|
| 1. BCCGCG | | | 1.0e-083 | 1.0e-083 | HINFP, Zfp161, GLIS2, TFDP1 |
| 2. CGCCG | | | 4.4e-077 | 6.8e-079 | Nr2f2, E2F2, Gata6, ELK1, E2F3 |
| 3. CGCCCS | | | 2.0e-066 | 5.1e-070 | Bcl6b, Sp4, Zfp410, KLF16, Egr1, KLF13, **SP1**, INSM1, **SMAD3**, etc. |
| 4. GCGSAS | | | 1.9e-063 | 2.2e-066 | HINFP, HINFP1, NRF1, Zfp161 ZEB1, FOXG1, YY2 |
| 5. CGCGM | | | 8.9e-062 | 1.1e-078 | Zfp161, ZBED1, ZBTB33, E2F2, E2F3, Tcfl5 |

**Figure S10: Confined migration increases chromatin accessibility at regions enriched with transcription factor motifs. (A)** Euler diagrams of "up" peaks in all three comparison groups: high versus low, medium versus low, and high versus medium collagen concentration samples. Numbers in each region represent the number of "up" peaks specific to the region. 5+21 (a total of 26) peaks are considered double-shared "up" peaks among the three comparison groups. **(B)** The signaling pathways or cellular functions of genes associated with double-shared "up" peaks. Green box: chromatin or gene silencing-related genes. Red box: cell migration or tumor invasion-related genes. SMAD3 is also a significant TF hit in the motif enrichment analysis. **(C-D)** qPCR analysis of HDAC3 and HP1α mRNA levels extracted from HT1080 cells migrating in low and high concentration collagen matrices. N.S., not significant, student's $t$ test with Welch's correction for unequal variances. Data are presented as mean ± SEM, from $n$ RNA lysates pooled from two biological replicates. **(E-F)** Quantification of HDAC3 and HP1α immunofluorescence labeling intensity in HT1080 cells migrating in microfluidic devices. All values are normalized to cells in the "before" region of control channels. $^{**}p < 0.01$, one-way ANOVA with Tukey's multiple comparison test. Data are presented as mean ± SEM, based on *n* cells (listed in each graph) pooled from three biological replicates. **(G)** Top 5 enriched DNA motifs in "up" peaks and their matching transcription factor (TF) hits. RC: reverse complement. E-value: enrichment *p*-value times the number of tested candidate motifs. Unerased E-value: E-value without erasing the motif sites previously found. Significant TF hits: TFs with matching binding motifs of $p$-value < 0.05. Underlined: SP1 and SMAD3, which are two promising TFs that may regulate "up" peaks-associated genes.

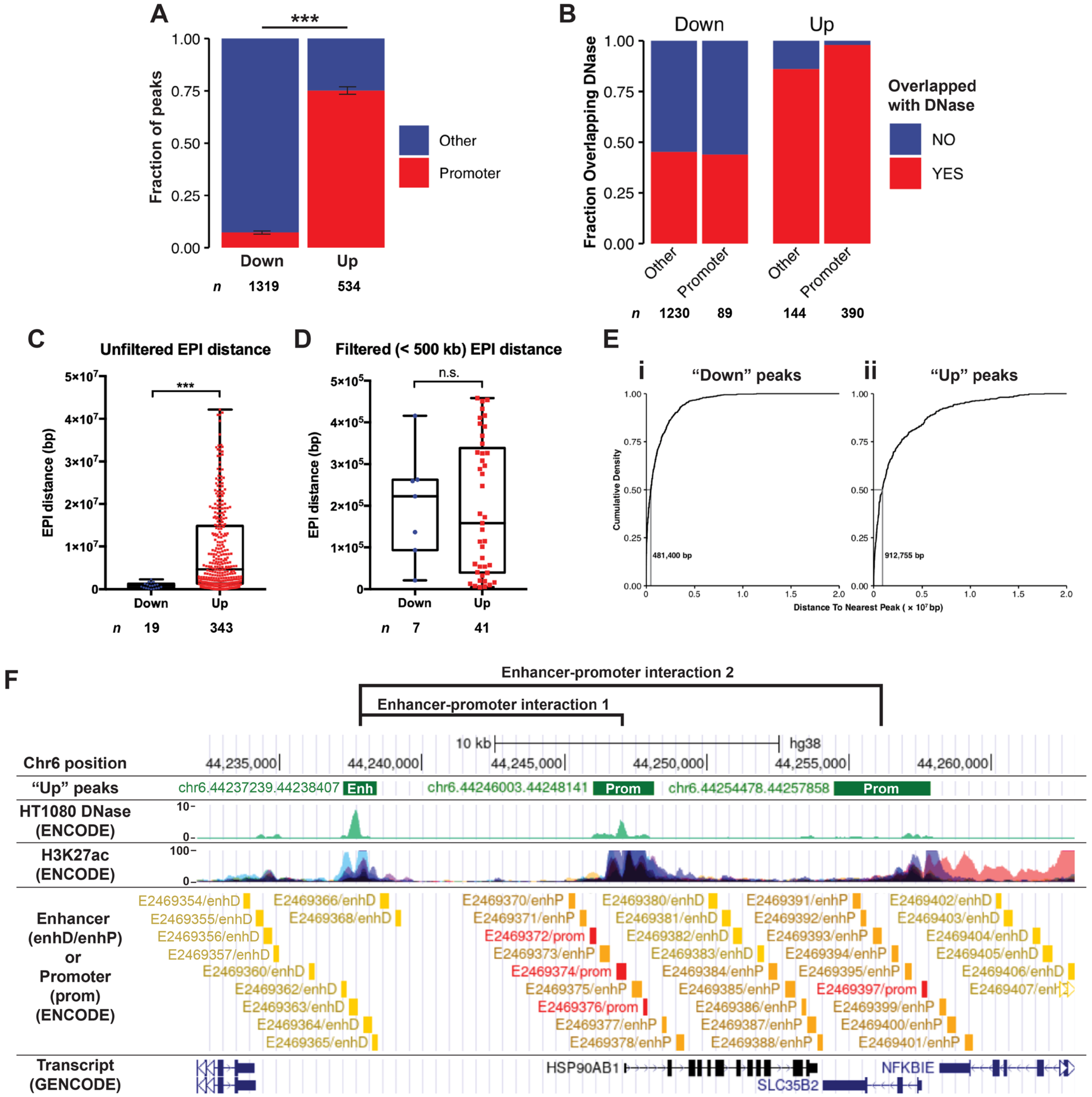


**Figure S11: Confined migration regulates co-accessibility of enhancer-promoter interactions. (A)** The fraction of promoter-associated peaks (within 1 kb upstream of TSS, in red) and non-promoter-associated peaks (other, in purple) within "down" and "up" peaks. ***$p < 0.001$, student's $t$ test with Welch's correction for unequal variances. Data are presented as mean ± SEM, based on $n$ DA peaks. **(B)** Fraction of promoter-associated (labeled in red) or non-promoter-associated (other, labeled in blue) peaks overlapping DNase accessibility data of HT1080 cells from ENCODE database. **(C)** Distribution of all enhancer-promoter interaction (EPI) distances within "down" and "up" peaks. ***$p < 0.001$, Wilcoxon Rank Sum test. Data are represented as scatter plots with median, box (1st and 3rd quartiles) and whiskers (minimum and maximum). **(D)** Distribution of filtered (< 500 kb cutoff) EPI distances within "down" and "up" peaks. N.s. not significant, Wilcoxon Rank Sum test ($p = 0.864$). Data are represented as scatter plots with median, box (1st and 3rd quartiles) and whiskers (minimum and maximum). **(E)** Empirical cumulative distribution plots of the distance to the nearest peak of (i) "down" peaks and (ii) "up" peaks, ranking from the smallest to the largest distance between two peaks. Y-axis: the cumulative distribution. X-axis: the distance to the nearest peak in $\times 10^7$ bp. The median distance is shown at x = 0.50. **(F)** Representative UCSC genome browser shots (http://genome.ucsc.edu) of two EPIs that exhibit co-accessibility changes (up-regulated in high collagen concentration sample). First (top) lane: The positions on chromosome 6. Second lane: Green blocks indicate the regions of “up” peaks. “Enh” indicates the enhancer-associated peak, and “Prom” indicates promoter-associated peaks. Third lane: DNase I hypersensitive assay data of HT1080 cells from ENCODE database, indicating accessible genomic regions. Fourth lane: H3K27ac ChIP-seq data from ENCODE database, indicating regulatory elements such as enhancers. Fifth lane: Enhancer (enhD for distal enhancer, and enhP for proximal enhancer) and promoter (prom) annotations from ENCODE database. Sixth (bottom) lane: Transcript annotation from GENCODE database.

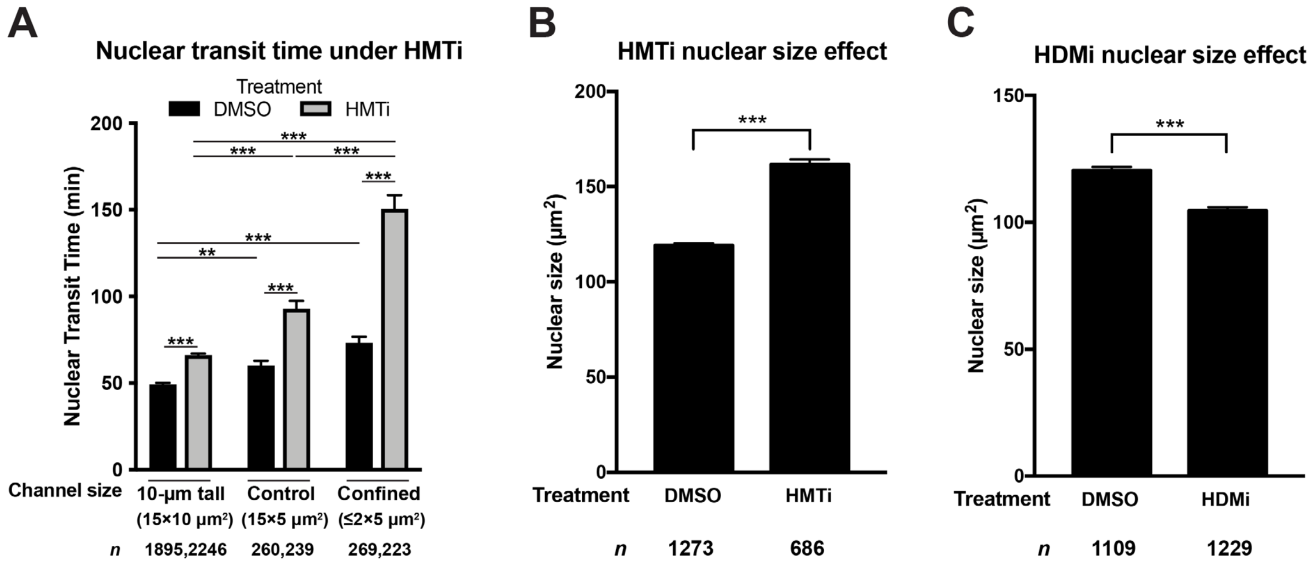


**Figure S12: The effect of histone modifying enzyme inhibitors on nuclear transit time and nuclear size.** **(A)** Quantification of HT1080 cells nuclear transit time through 10-μm tall (15×10 μm$^2$), control (15×5 μm$^2$) or confined (≤2×5 μm$^2$) channels, treated with vehicle (DMSO) or HMTi. **$p < 0.01$, ***$p < 0.001$, Kruskal-Wallis H test with Dunn's multiple comparison test. **(B)** Quantification of HT1080 cells nuclear size under DMSO (vehicle) or HMTi treatment. ***$p < 0.001$, student's *t* test with Welch's correction for unequal variances. **(C)** Quantification of HT1080 cells nuclear size under DMSO (vehicle) or HDMi treatment. ***$p < 0.001$, student's *t* test with Welch's correction for unequal variances. Data are presented as mean ± SEM, based on *n* cells (listed in each graph) pooled from three biological replicates.

**Supplemental Tables**

**Table S1: Representative genes associated with up- or down- regulated DA peaks in high concentration collagen samples.** FC: fold change. Padj: adjusted *p*-value.

| DA direction | Log2FC | padj | Gene name |
|---|---|---|---|
| Up | 0.543 | 3.071E-03 | *TRIM44* |
| Up | 0.213 | 8.277E-02 | *HDAC3* |
| Up | 0.204 | 3.199E-02 | *CBX5 (HP1α)* |
| Down | -1.193 | 1.374E-19 | *TERT* |
| Down | -1.406 | 1.342E-08 | *COL23A1* |
| Down | -1.469 | 6.419E-09 | *MUC2* |

**Table S2: Chromatin/gene silencing or other chromatin-related genes associated with up-regulated DA peaks in high concentration collagen samples.** FC: fold change. Padj: adjusted *p*-value. Histone protien family names are highlighted with underline. *Genes encoding critical proteins involved in heterochromatin formation and maintenance. †SMAD3 is also a significant hit in motif analysis.

| Log2FC | padj | Gene name | GO: Chromatin silencing or Gene silencing | GO: Telomeric region |
|---|---|---|---|---|
| 0.577 | 7.123E-03 | *ANP32B* | | |
| 0.453 | 1.402E-03 | *MOV10* | V | |
| 0.426 | 4.364E-02 | *ACIN1* | | |
| 0.406 | 6.468E-02 | *POLR2K* | | |
| 0.399 | 6.708E-06 | *HIST1H2AC* | V | |
| 0.385 | 2.747E-05 | *HIST3H2A* | V | |
| 0.373 | 9.991E-04 | *HIST2H4A* | V | V |
| 0.373 | 9.991E-04 | *HIST2H4B* | V | V |
| 0.360 | 9.399E-02 | *SMAD3*† | | |
| 0.338 | 7.605E-04 | *HIST1H4A* | V | V |
| 0.326 | 2.960E-04 | *HIST2H2AC* | V | |
| 0.315 | 5.005E-02 | *DDX21* | | |
| 0.308 | 2.489E-03 | *H3F3B* | V | V |
| 0.305 | 7.140E-02 | *HIST1H2AJ* | V | |
| 0.297 | 3.410E-02 | *HIST1H2AH* | V | |
| 0.285 | 6.113E-02 | *AGO3* | V | |
| 0.279 | 3.121E-02 | *HIST1H2AB* | V | V |
| 0.272 | 7.048E-02 | *FEN1* | | V |
| 0.258 | 9.595E-02 | *HIST1H2AI* | V | V |
| 0.256 | 8.714E-02 | *ORC2* | | V |
| 0.253 | 8.406E-02 | *THOC5* | | V |
| 0.249 | 3.500E-02 | *OIP5* | | |
| 0.248 | 9.811E-02 | *EID3* | | V |
| 0.246 | 3.049E-02 | *HIST1H2BD* | | |
| 0.245 | 8.069E-02 | *HAT1** | V | V |
| 0.235 | 4.155E-02 | *H2AFX* | V | V |
| 0.230 | 3.683E-02 | *HIST1H2AG* | V | |
| 0.230 | 6.372E-02 | *CENPO* | | |
| 0.226 | 8.924E-02 | *HIST2H3A* | V | |
| 0.215 | 9.811E-02 | *NCAPD2* | | |
| 0.213 | 8.277E-02 | *HDAC3** | | |
| 0.204 | 3.199E-02 | *CBX5/HP1α** | | V |

**Table S3: DNA damage checkpoint genes associated with up-regulated DA peaks in high concentration collagen samples.** FC: fold change. Padj: adjusted *p*-value.

| Log2FC | padj | Gene name |
|---|---|---|
| 0.362 | 4.317E-03 | *UBC* |
| 0.346 | 9.811E-02 | *DTL* |
| 0.342 | 9.746E-02 | *TRIAP1* |
| 0.329 | 8.837E-02 | *GML* |
| 0.323 | 1.168E-02 | *PRPF19* |
| 0.319 | 2.434E-02 | *ZNF385A* |
| 0.317 | 3.268E-02 | *ATRIP* |
| 0.310 | 2.869E-02 | *MDM2* |
| 0.291 | 5.580E-02 | *PTPN11* |
| 0.289 | 5.177E-02 | *MDM4* |
| 0.289 | 7.325E-02 | *FANCI* |
| 0.276 | 5.722E-02 | *SYF2* |
| 0.258 | 5.080E-02 | *MAPK14* |
| 0.235 | 4.155E-02 | *H2AFX* |

**Table S4: Cell cycle chekpoint genes associated with up-regulated DA peaks in high concentration collagen samples.** FC: fold change. Padj: adjusted *p*-value.

| Log2FC | padj | Gene name |
|---|---|---|
| 0.369 | 7.233E-03 | *RB1* |
| 0.300 | 3.116E-02 | *AURKB* |
| 0.250 | 3.575E-02 | *E2F7* |
| 0.239 | 7.223E-02 | *MAD2L1* |
| 0.221 | 9.978E-02 | *BUB3* |
| 0.212 | 9.444E-02 | *CDC45* |

**Table S5: Gene Ontology (GO) Cellular Component terms enrichment of genes associated with up-regulated DA peaks in high concentration collagen samples.** *Chromatin sturcture or gene silencing-related GO terms.

| GO term | Binominal rank | Binominal *p*-value |
|---|---|---|
| DNA packaging complex* | 1 | 6.5383e-14 |
| Nucleosome* | 2 | 1.5848e-13 |
| Protein-DNA complex* | 8 | 8.2620e-8 |
| Nuclear nucleosome* | 11 | 1.7191e-5 |
| Chromosome, telomeric region* | 13 | 5.8580e-5 |
| Nuclear chromosome, telomeric region* | 14 | 6.0764e-5 |
| Chromosomal region* | 15 | 8.0535e-5 |

**Table S6: List of primers used for qPCR.**

| Gene (all human) | Primer sequences (5' to 3') |
|---|---|
| *β-actin* | Fwd: CCCCGCGAGCACAGAG<br>Rev: ATCATCCATGGTGAGCTGGC |
| *GAPDH* | Fwd: TGCGTCGCCAGCCGAG<br>Rev: AGTTAAAAGCAGCCCTGGTGA |
| *18S* | Fwd: GGCCCTGTAATTGGAATGAGTC<br>Rev: CCAAGATCCAACTACGAGCTT |
| *HDAC3* | Fwd: GGCAACTTCCACTACGGAGC<br>Rev: GGCCTGGTATGGCTTGAAGA |
| *CBX5/HP1α* | Fwd: CTCTCAATCCCGGGGACCT<br>Rev: CAGCTGTCCGCTTGGTTTTC |

**Table S7: Thermocycler settings used for qPCR.**

| Step | Cycles | Temperature (°C) | Duration |
|---|---|---|---|
| Pre-incubation | 1 | 95 | 10 min |
| Amplification | 50 | 95<br>60<br>72 | 15 sec<br>30 sec<br>30 sec |
| Melting curve | 1 | 95<br>50<br>97 | 5 sec<br>1 min<br>continuous |
| Cooling | 1 | 40 | 30 sec |

**Supplementary Videos**

**Video S1: Nuclear transit through confined constrictions induces GFP-HP1α enrichment formation in an HT1080 cell.** Green (left panel) and inverted grayscale (right panel): GFP-HP1α. White (left panel) and blue (right panel) arrowheads: GFP-HP1α enrichment. Scale bar: 10 μm.

**Video S2: Long-term time-lapse tracking of confined migration followed by immunofluorescence staining of histone marks in HT1080 cells.** Green: NLS-GFP. Arrowhead: tracking of the cell of interest ("Cell A" in Supplementary Fig. 5b). Red: H3K27me3 staining. Cyan: H3K9ac staining. Scale bar: 50 μm.

**Video S3: Nuclear transit through confined constrictions does not change the nucleoplasmic-to-cytoplasmic ratio of GFP-HDAC3 in an HT1080 cell.** Green (left panel) and inverted grayscale (right panel): GFP-HDAC3. Red: SPY555-DNA. Cyan lines: timing and threshold of nuclear entry and exit of the constriction. Scale bar: 15 μm.

**Video S4: Nuclear transit of an HT1080 cell through control (15×5 $\mu m^2$) constrictions under the vehicle control (DMSO).** Green: NLS-GFP. Red: H2B-mCherry. Cyan lines: timing and threshold of nuclear entry and exit of the constriction. Scale bar: 10 μm.

**Video S5: Nuclear transit of an HT1080 cell through control (15×5 $\mu m^2$) constrictions under a histone methyltransferase inhibitor (HMTi).** Green: NLS-GFP. Red: H2B-mCherry. Cyan lines: timing and threshold of nuclear entry and exit of the constriction. Scale bar: 10 μm.

**Video S6: Nuclear transit of HT1080 cells through confined (≤2×5 $\mu m^2$) constrictions under the vehicle control (DMSO).** Green: NLS-GFP. Red: H2B-mCherry. Cyan and magenta lines: timing and threshold of nuclear entry and exit of the constriction. Scale bar: 10 μm.

**Video S7: Nuclear transit of HT1080 cells through confined (≤2×5 $\mu m^2$) constrictions under a histone methyltransferase inhibitor (HMTi).** Green: NLS-GFP. Red: H2B-mCherry. Cyan and magenta lines: timing and threshold of nuclear entry and exit of the constrictions. Scale bar: 10 μm.

**Video S8: Nuclear transit of an HT1080 cell through confined (≤2×5 $\mu m^2$) constrictions under mild hypothermia and the vehicle control (DMSO).** Green: NLS-GFP. Red: H2B-mCherry. Cyan lines: timing and threshold of nuclear entry and exit of the constriction. Scale bar: 10 μm.

**Video S9: Nuclear transit of an HT1080 cell through confined (≤2×5 $\mu m^2$) constrictions under mild hypothermia and a histone demethylase inhibitor (HDMi).** Green: NLS-GFP. Red: H2B-mCherry. Cyan lines: timing and threshold of nuclear entry and exit of the constriction. Scale bar: 10 μm.
